# Predicting tumor immune microenvironment and checkpoint therapy response of head & neck cancer patients from blood immune single-cell transcriptomics

**DOI:** 10.1101/2023.01.17.524455

**Authors:** Yingying Cao, Tiangen Chang, Fiorella Schischlik, Kun Wang, Sanju Sinha, Sridhar Hannenhalli, Peng Jiang, Eytan Ruppin

**Affiliations:** Cancer Data Science Laboratory, Center for Cancer Research, National Cancer Institute (NCI), National Institutes of Health (NIH), Bethesda, MD, USA 20892; Boehringer Ingelheim RCV Gmbh & Co KG, Vienna, Austria

## Abstract

The immune state of tumor microenvironment is crucial for determining immunotherapy response but is not readily accessible. Here we investigate if we can infer the tumor immune state from the blood and further predict immunotherapy response. First, we analyze a dataset of head and neck squamous cell carcinoma (HNSCC) patients with matched scRNA-Seq of peripheral blood mononuclear cells (PBMCs) and tumor tissues. We find that the tumor immune cell fractions of different immune cell types and many of the genes they express can be inferred from the matched PBMC scRNA-Seq. Second, analyzing another HNSCC dataset with PBMC scRNA-Seq and immunotherapy response, we find that the inferred ratio between tumor memory B and regulatory T cell fractions is predictive of immunotherapy response and is superior to the well-established cytolytic and exhausted T-cell signatures. Overall, these results showcase the potential of scRNA-Seq liquid biopsies in cancer immunotherapy, calling for their larger-scale testing.

**Significance:** This head and neck cancer study demonstrates the potential of using blood single-cell transcriptomics to (1) infer the tumor immune status and (2) predict immunotherapy response from the tumor immune status inferred from blood. These results showcase the potential of single-cell transcriptomics liquid biopsies for further advancing personalized cancer immunotherapy.

## Introduction

The immune status of tumor microenvironment (TME) plays a critical role in determining the progression of cancer and the efficiency of immunotherapy. Two important determinants of the immune status in the TME are the composition and gene expression of different immune cell types. In terms of cell abundance, a strong infiltration of CD8+ T cells is associated with favorable patient prognosis [1, 2], while a high infiltration of regulatory T cells (Treg) correlates with poor prognosis in many solid tumors [3, 4]. A high CD8+/Treg ratio is associated with favorable prognosis in multiple tumor types [5, 6]. Additionally, tumor-infiltrating neutrophils, lymphocytes, and neutrophils-to-lymphocytes ratio have been found to be important markers of patient survival [7]. Beyond cell abundance, the gene expression in the TME has been shown to be predictive of cancer immune checkpoint blockade (ICB) response, for example, by identifying gene expression signatures of T cell dysfunction and exclusion [8] or by pairwise transcriptomic relations between immune checkpoint genes (ICGs) [9].

The rapidly evolving single-cell sequencing technology now offers an unprecedented opportunity to analyze cell composition and gene expression of tumor cells as well as immune cells. For example, this technology can be used to study genomic and transcriptomic heterogeneity of tumors [10], construct a tumor immune cell atlas [11], chart cell-cell communication in the TME [12, 13], understand genomic and transcriptomic dynamics during cancer progression [14-16] and track tumor evolution [17]. In the clinic, ideally, one would like to perform multiple tumor biopsies and employ single-cell transcriptomics to study the dynamics of the TME. However, for many tumors, such invasive procedures are prohibitive. In this study, we aim to determine the extent to which we can learn about the state of the TME from single-cell transcriptomics of immune cells in the blood.

Liquid biopsies have greatly advanced the field of clinical oncology and have become a focus of ongoing research for precision diagnosis and personalized cancer treatment. Current efforts have been focused on studying cell-free circulating tumor DNA (ctDNA), circulating tumor cells (CTCs) and methylation signatures (see [18] for a review). Here our focus is different and complementary, namely, to study single cell transcriptomics of immune cells in the blood to learn about the immune status of the TME. Previous related studies have aimed to predict normal tissue gene expression from the *bulk* expression of white blood cells (WBC). For example, analyzing matched whole blood/lung gene expression data from the Genotype-Tissue Expression (GTEx) project, a generalized linear regression model was shown to predict the expression levels of ∼18% of the genes in the lung from the blood [19]. Using a Bayesian ridge regression-based method, the expression levels of 20%∼60% genes in 16 tissues could be predicted from the blood [20]. Recent research has also shown that whole blood transcriptomes can predict tissue-specific expression levels for ∼60% of the genes on average across 32 tissues, and the inferred tissue-specific expression from the blood transcriptome can predict the disease state in some disorders [21].

Encouraged by our previous success with predicting tissue-specific expression from an individual’s blood bulk transcriptome [21], in this study we first aim to learn to what extent can one use single cell blood transcriptome to learn about the cellular composition and gene expression of various immune cell types within an individual’s TME. Second, based on that, we aim to learn if we can use the inferred TME immune state to predict ICB response, and furthermore, if that leads to more accurate predictions than predictors that are built directly on the blood information without inferring the TME immune state.

To this end, we first analyzed a dataset of tumor-blood matched single-cell RNA sequencing (scRNA-Seq) data for head and neck squamous cell carcinoma (HNSCC) from 26 patients. We hypothesized that immune cells in the peripheral blood mononuclear cells (PBMCs) bear information on the immune state in the TME, due to ongoing circulation of both immune cells and immune signaling molecules like cytokines between the blood and the TME. Indeed, many previous studies have shown that the number and function of both innate and adaptive immune cells in the blood are altered in cancer patients [22-24]. Additionally, it has been found that the dynamics of circulating immune cell phenotypes reflect interactions between tumor and immune cells during immunotherapy [25]. Furthermore, numerous studies have also reported that changes in blood immune cell composition, particularly in neutrophils-to-lymphocytes ratio, are associated with poor survival outcomes and response to immunotherapy in many types of solid tumors [26]. Indeed, we find that the immune cell fractions (ICFs) of major immune cell types in the TME can be predicted from PBMC scRNA-Seq as they are correlated with ICFs of certain cell types in the latter. ICF is a proxy for cell abundance and represents the fraction of cells of a given immune cell type out of the total number of all viable CD45+ immune cells in the same sample which includes almost all immunological and hematological cells except for mature erythrocytes and platelets [27, 28]. Additionally, we find that in different immune cell types in the TME the expression levels of 17-47% expressed genes can be predicted from the patient’s blood.

We then analyzed a second data set of HNSCC patients including both patient pre-treatment blood scRNA-Seq data and ICB response information. We set to study if we can predict ICB response based on patients’ PBMC scRNA-Seq data. We find that the well-established exhausted T-cell signature in the TME can be inferred from the blood and the inferred exhausted T-cell score can further predict immunotherapy response in HNSCC. In addition, we identify a new immune signature named ICFR*, which reflects the blood-inferred TME ICF ratio of (B_memory_ - T_reg_) / (B_memory_ + T_reg_). The inferred ICFR* scores in the TME are the most predictive of patients’ response to ICB treatment both in terms of tumor RECIST criteria and overall response in this dataset, and importantly, are further validated in an independent large scale bulk expression HNSCC dataset.

## Results

### The data analyzed and analysis overview

We obtained the single cell transcriptome data from [29], where all viable CD45+ immune cells were measured for matched PBMC samples and primary tumor tissue samples from 18 HPV– and 8 HPV+ immunotherapy treatment naïve HNSCC patients. Immune cell identities were annotated using *CellTypist*, an automated cell classification tool designed by [30]. The resulting cell identities were then merged into 12 major immune cell types such that each cell type in tumor tissues had an average immune cell fraction (ICF) > 1% (**Methods**). ICF is defined as the fraction of cells of a given immune cell type among all CD45+ cells in the same sample. Specifically, the 12 immune cell types identified in this study are cycling T cells, cytotoxic T cells, dendritic cells, germinal center B cells, helper T cells, macrophages, memory B cells, monocytes, naïve B cells, natural killer (NK) cells, plasma cells, and regulatory T cells (**Fig. 1A**). Throughout this study we quantify cell abundances in term of ICF.

**Figure 1.**
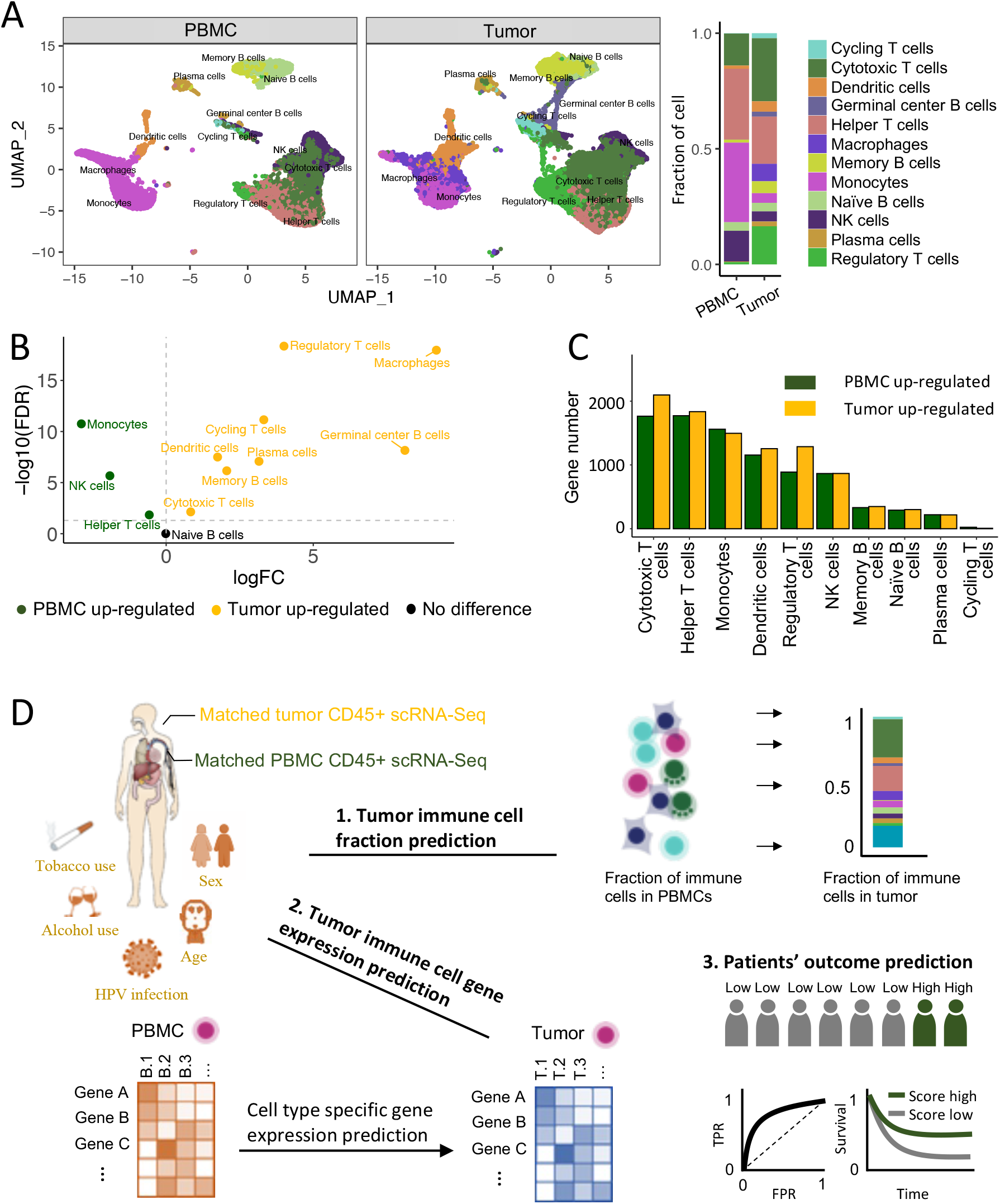
Data basic properties and analysis overview. **A**. Distribution, annotation, and cell fractions of 12 major immune cell types in matched tumor-PBMC CD45+ scRNA-Seq data from 26 treatment-naïve HNSCC patients. **B**. A volcano plot showing differential fractions of abundance of 12 major immune cell types between tumor tissues and PBMCs. **C**. Significantly differentially expressed genes in 12 annotated immune cell types from tumor tissues and PBMCs. **D**. An overview of the study: Firstly, **(1)** the ICFs andthe gene expression levels of major immune cell types in the TME are predicted from the blood scRNA-Seq data. Then, **(3)** immune signatures are identified from immune information in the blood and in the TME (which is predicted from the blood) to predict patients’ ICB response.

We first compared the similarity and difference of the immune microenvironment between the TME and the blood of HNSCC patients. We observed that the ICFs of all immune cell types, except for naïve B cells, are significantly different between tissue samples and matched PBMC samples.

Specifically, the ICFs of cycling T cells, cytotoxic T cells, dendritic cells, germinal center B cells, macrophages, memory B cells, plasma cells, and regulatory T cells are higher in the TME, while the ICFs of helper T cells, monocytes, and NK cells are lower in the TME (**Fig. 1A, B**).

When comparing the same immune cell types between the blood and the TME, we found that the expression levels of hundreds to thousands of genes are significantly altered. Across different immune cell types, there are up to 1772 genes that are significantly upregulated in PBMC samples and up to 2097 genes that are significantly upregulated in the tumor (log2 fold change > 0, FDR < 0.05; **Fig. 1C**).

Specifically, genes that are upregulated in the TME are enriched in specific immune response functions (**Supplementary Fig. 1**). Our functional analysis revealed that genes upregulated in PBMCs were more enriched in basic housekeeping biological functions such as “rRNA processing” and “cytoplasmic translation”, while genes upregulated in TME were more enriched in anti-tumor immune functions such as “T cell activation” and “lymphocyte mediated immunity” (**Supplementary Fig. 2**).

Next, we evaluated the extent to which we could predict the immune status in the TME from the blood using a machine learning based framework (**Fig. 1D**). Briefly, we first predicted the ICFs of 11 out of 12 major immune cell types in the TME using the cell types’ ICFs in the matching blood sample and additional clinical information, such as HPV infection status, alcohol and tobacco use, age, and sex.

Germinal center B cells were excluded from this analysis as they were found only present in HPV+ tumor tissue samples in this study. Second, we predicted the expression level of each gene in each immune cell type in the TME based on its expression level in the corresponding cell type in the matching blood sample along with the patient’s clinical information. Finally, using the immune status information learned from the blood, we identified immune signatures that are predictive of HNSCC patients’ ICB response.

### The immune cell fractions (ICFs) of all cell types in the TME can be predicted from the blood

We first explored the correlation between ICFs in the TME and those in the blood. To reduce the impact of potential sampling bias, correlations between ICFs were calculated with bootstrapping by randomly re-sampling the 25 patient samples for 1000 replications (note that sample 5 was excluded from the 26 patient samples as an outlier; **Supplementary Fig. 3**). For different cell types in the TME, the highest correlations with cell types in the blood were very different, with the Pearson correlations ranging from 0.14 to 0.69 (**Fig. 2A**). More intriguingly, for six out of 11 cell types in the TME, the most correlated cell types in the blood are different from themselves (**Fig. 2A**). Among these six cell types, two are not present in the blood and hence matching cells could not have been found (cycling T cells and macrophages); two are moderately positively correlated with themselves between the TME and the blood (plasma cells and regulatory T cells); however, the ICFs of two cell-types in the blood and in the tumor samples are not correlated at all (memory B cells and NK cells), suggesting that they may play different functional roles in the TME versus in the blood, or that their recycling rates between the tumor and the blood are low.

**Figure 2.**
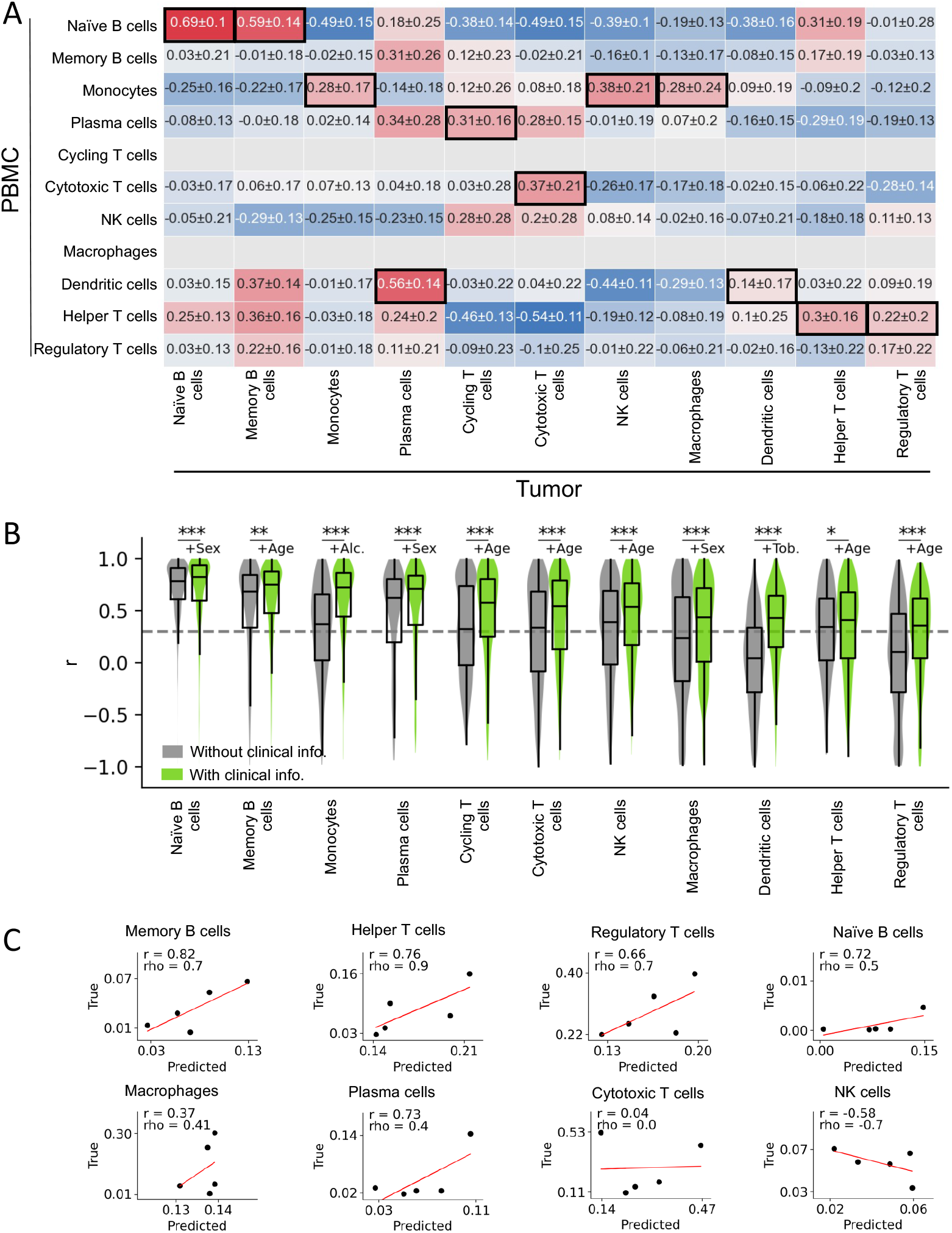
The ICFs of major immune cell types in the TME can be predicted from the blood. **A**. The correlation matrix of ICFs between the TME and the blood. For immune cell types in the TME (columns), the cell types in the blood (rows) with the highest correlations are marked in black boxes. Pearson correlation values shown are mean ± s.d. over 1000-replicate bootstrapping. **B**. Distribution of correlations between the model-predicted ICFs and the true TME ICFs are shown on the test sets across 1000 replicates. **C**. The correlation between the model-predicted TME ICF and the measured TME ICF for each cell type on a small, independent validation dataset (n = 5). Cycling T cells are not studied as they are not present in all tumor tissue samples; monocytes and dendritic cells are excluded due as their pertaining predictive clinical information is missing. Abbreviations: Alc., alcohol; Tob., tobacco; r denotes the Pearson correlation coefficient and rho denotes the Spearman correlation coefficient. Statistical notation: ***, FDR < 0.001; **, FDR < 0.01; *, FDR < 0.05. FDRs are calculated using Benjamini-Hochberg correction.

We then aimed to improve the prediction of ICFs in the tumor from the blood ICFs beyond the basic correlations denoted above by building machine learning predictors. Due to the small size of the training data, we focused on building very compact prediction models to minimize the risk of overfitting as much as possible. Specifically, for each cell type in the TME, a 1-variable linear regression model was built by using the ICF of its most correlated cell type in the blood (as the only variable), and five 2-variable linear regression models were built using the ICF of its most predictive cell type in the blood (as the first variable) and one of the five clinical variables (as the second variable). The most predictive model among those six models was chosen as the final model for predicting that cell-type’s ICF. To further overcome overfitting, we randomly split the data into training and test sets (80%: 20%) for 1000 replicates and performed model training and testing on the 1000 folds for each cell type (**Methods**). As shown in **Fig. 2B**, the ICFs of all 11 immune cell types in the TME were predictable, with mean Pearson correlation coefficients between the predicted and the true ICFs greater than 0.3 (r > 0.3), mean normalized mean absolute error less than 1 (NMAE < 1) and a Benjamini-Hochberg adjusted p-value < 0.05 (FDR < 0.05; **Fig. 2B**). Notably, clinical information played a statistically significant role in increasing the predictability of the ICFs for all immune cell types, compared to the 1-variable baseline models (**Fig. 2B**).

As these models were generated and tested via a standard cross validation procedure on the training set, we further tested them (without any further changes) on a smaller, independent, recently published external dataset (n = 5; [31]). Despite the very limited number of matched samples, quite strikingly, 6 out of 8 cell types whose ICFs were predictable in the original datasets were also predictable in this test set (with r > 0.3). These included the memory B cells, helper T cells, regulatory T cells, naïve B cells, macrophages, and plasma cells. Notably, memory B cells, helper T cells, and regulatory T cells were well-predicted with both r > 0.6 and rho (the Spearman correlation coefficient) > 0.6 (**Fig. 2C**).

### The expression levels of 17-47% of the genes that are expressed in specific immune cell types in the TME can be predicted from the blood

Similarly, we first examined the correlation between gene expression levels of immune cell types in the TME and those in the blood. As expected, we found that the gene expression levels of all cell types in the TME were most highly correlated with those of the same cell types in the blood, except for cycling T cells and macrophages, which were not found in the blood (**Supplementary Fig. 4**).

To predict gene expression levels of each immune cell type in the TME, we used a similar model construction strategy. For each gene in each cell type, a 1-variable linear regression model was built by using the expression pattern of the gene itself in the corresponding cell type in the blood (as the only variable), and five 2-variable linear regression models were constructed using the expression level of the gene itself in the corresponding cell type in the blood (as the first variable) and one of the five clinical variables (as the second variable). The model that had the strongest mean predictive power on the test sets over 1000 replicates was chosen as the final model. (**Methods**).

As observed in the prediction of for ICF, clinical information plays a significant role in the prediction of gene expression as well (**Fig. 3A**). Notably, various clinical variables appear to have varying levels of significance across different cell types. HPV infection status is the most crucial factor in predicting the expression of dendritic cells and all T cells, tobacco use is the most crucial factor for innate immune cells (i.e., monocytes, macrophages, and NK cells), and alcohol use is the most crucial factor for plasma cells and all B cells (**Fig. 3B**).

**Figure 3.**
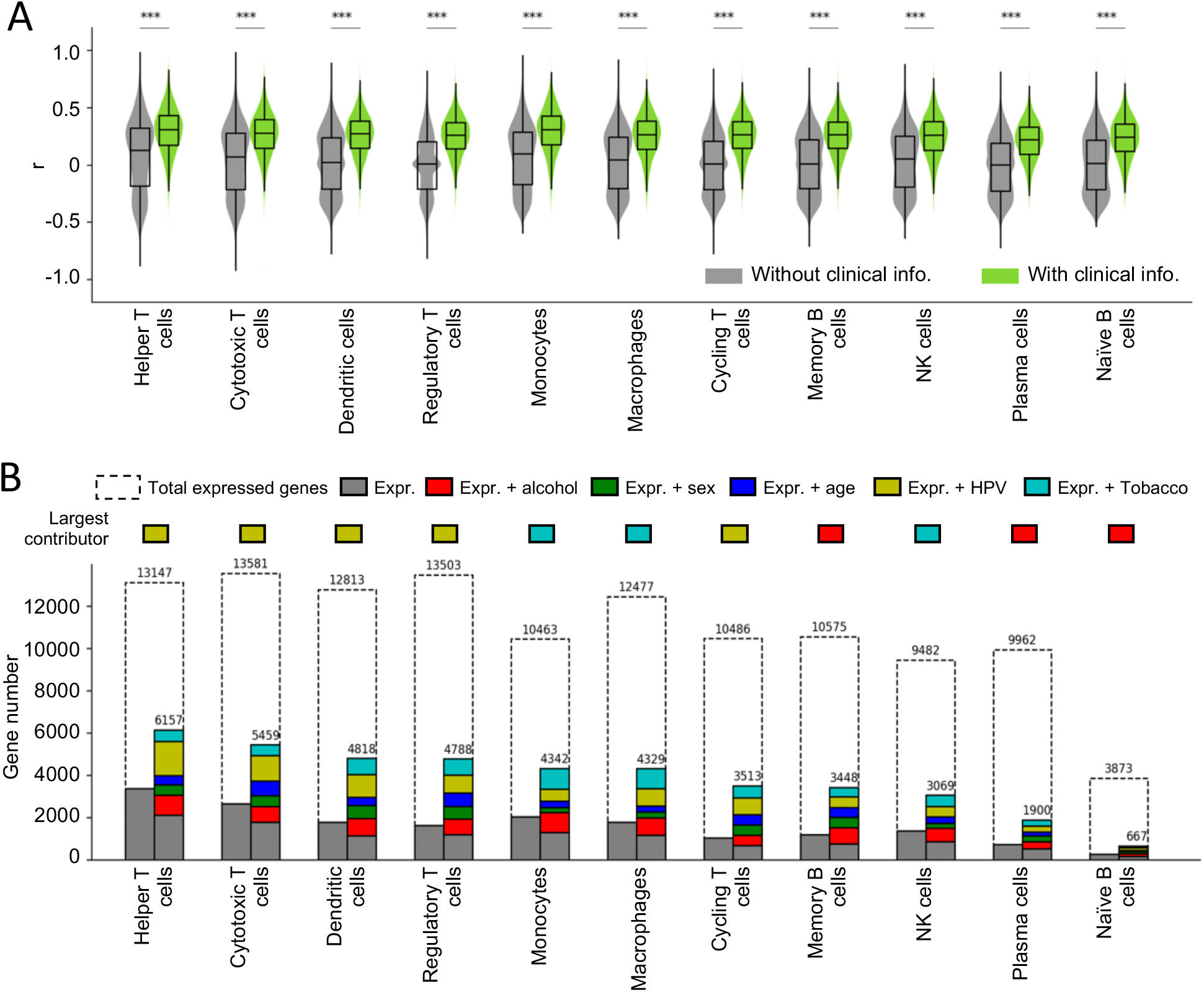
The expression levels of 17-47% of the genes expressed in TME immune cells can be predicted from the blood. **A**. The distribution of correlations between the predicted and true expression values of all expressed genes in each immune cell type in the TME, with and without clinical information. **B**. The number of expressed genes and predictable genes in each cell type, with and without the consideration of clinical variables. The grey bars represent the number of genes that can be predicted using expression alone, while the colored bars indicate the additional number of genes that can be predicted by incorporating clinical variables (see the corresponding-colored labels). Statistical notation: ***, FDR < 0.001; **, FDR < 0.01; *, FDR < 0.05. The FDRs are calculated using Benjamini-Hochberg correction method.

Overall, the expression levels of 17-47% of the genes expressed in immune cells in the TME can be predicted from the blood (mean r > 0.3 and mean NMAE < 1 with FDR < 0.05 on the test sets over 1000 replicates; **Fig. 3B**). Notably, dendritic cells and helper, cytotoxic, and regulatory T cells have the highest number of predictable genes (> 4700), while plasma cells and naïve B cells have the least (< 2000 and < 700 genes respectively). The detailed lists of predictable genes in different immune cell types can be found in **Supplementary Material SM1**. Furthermore, our findings indicate that 22-57% of the significantly differentially expressed genes between the TME and the blood (**Fig. 1C**) are predictable across different immune cell types (**Supplementary Fig. 5**).

Functional gene ontology (GO) enrichment analyses of the predictable genes in different immune cell types in the TME reveals a number of common immune response-related GO terms shared across different immune cell types, such as “adaptive immune response”, “leukocyte cell-cell adhesion”, “positive regulation of T cell activation”, “response to virus”, “T cell activation”, “antigen processing and presentation” and “Th17 cell differentiation” (**Supplementary Table 2**). Notably, genes that are most accurately predicted by the HPV infection status information are enriched in virus infection related terms. For example, in helper T cells, these genes are enriched in “DNA repair”, “HIV infection”, and “HIV life cycle”; and in regulatory T cell, these genes are enriched in “influenza infection” (**Supplementary Fig. 6**).

### Exhausted T-cell signature in the TME can be inferred from the blood and the inferred exhausted T-cell score can further predict immunotherapy response in HNSCC

Given the predictability of ICFs and gene expression levels of different immune cell types in the TME, we were interested to test if we could predict the scores of some well-established immune signatures in the TME from the PBMC scRNA-Seq data. Specifically, we checked the predictability of three pan-cancer T cell immune signatures for immunotherapy including the exhausted T-cell signature (average expression of PDCD1, CTLA4, LAG3, HAVCR2 and TIGIT in cytotoxic T cells) [32], the cytolytic T-cell signature (average expression of GZMA and PRF1 in cytotoxic T cells) [33], and CXCL13 expression in cytotoxic T cells [34]. This investigation was performed by analyzing a dataset of HNSCC patients [31], including both patient pre-treatment blood scRNA-Seq data and ICB response information.

Firstly, we found that the exhausted T-cell signature in the TME could be predicted from blood cytotoxic T cell gene expression levels in both the training dataset of [29] (r = 0.56; rho = 0.61) and the validation dataset of [31] (r = 0.46; rho = 0.40) (**Fig. 4A**). Moreover, the inferred exhausted T-cell signature in the TME was significantly negatively associated with patients’ ICB RECIST response (AUC = 0.88, p = 0.01; **Fig. 4B**), as one would expect. However, the exhausted T-cell signature was not significantly associated with patients’ survival after ICB treatment (**Fig. 4C**). This could be attributed to the relatively short follow-up time of 36 months, which might not have been sufficient to observe differences in survival among patients. It is worth noting that the exhausted T-cell signature score in the blood on its own is not correlated with that in the TME (**Fig. 4D**) and is not associated with ICB response or survival (**Fig. 4E, F**), highlighting the importance of predicting ICB response through the two-step approach. Finally, the cytolytic T-cell and CXCL13 signatures in the TME were less predictable from the blood than the exhuastion siganture (**Supplementary Fig. 7A, D)**. As a result, the predicted scores for these signatures had low correlations with ICB response and survival (**Supplementary Fig. 7B, C, E, F**).

**Figure 4.**
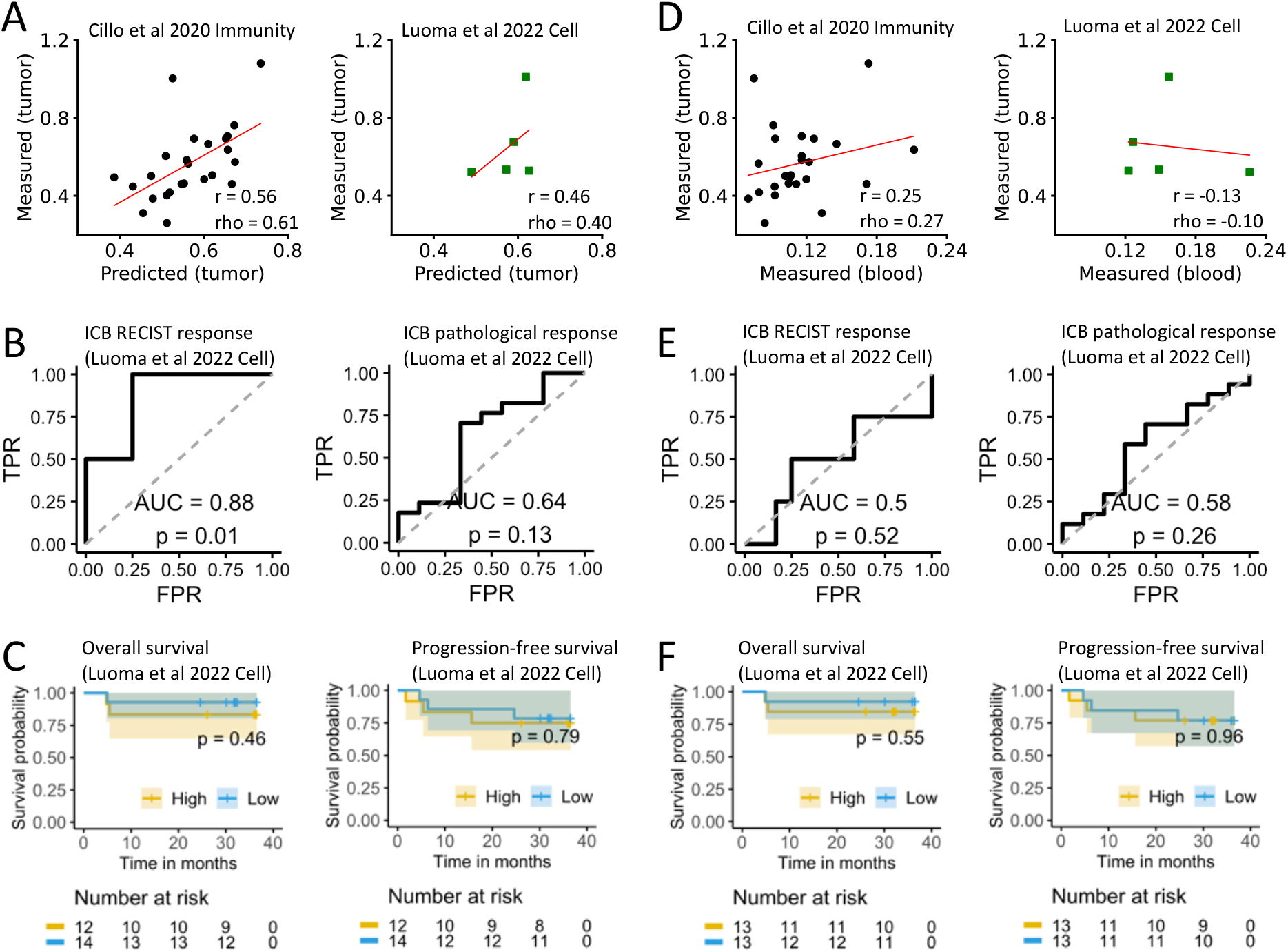
Exhausted T-cell signature in the TME can be inferred from the blood and the inferred exhausted T-cell score can further predict immunotherapy response in HNSCC. **A**. Correlation between the predicted and the measured exhausted T-cell signature scores in TME in the matched blood/tumor training dataset (n = 25) and validation dataset (n = 5). **B**. ROC curve for predicting ICB RECIST response (n = 16) and pathological response (n = 26) by the predicted exhausted T-cell signature scores. **C**. Overall survival and progression-free survival analyses of ICB-treated patients in exhausted-high (exhausted score > quantile 50%) versus exhausted-low (exhausted score ≤ quantile 50%) tumor groups using the predicted exhausted T-cell signature scores. Panels **D, E, F** are the same to A, B, C, respectively, except that the predicted exhausted T-cell signature scores in the TME are replaced by the measured exhausted T-cell signature scores in the blood. Abbreviations: r, Pearson correlation coefficient; rho, Spearman correlation coefficient; TPR, true positive rate; FPR, false positive rate.

### Prediction of HNSCC immunotherapy response via a newly identified *TME immune signature, ICFR\**

We next examined whether we could identify new blood-based immune signatures that can help predict HNSCC patients’ ICB response. To this end, we tested two different strategies. Strategy 1 was a direct approach, in which we predicted ICB response based directly on the immune information in the blood. Strategy 2 was an indirect, two-step approach, in which we first identified predictable immune information in the TME from the blood (**Figs. 2, 3**), and then used this information to predict ICB response.

To achieve our goal, we comprehensively examined three categories of immune information in both the blood (for “Strategy 1” candidates) and the corresponding predictable immune information in the TME (for “Strategy 2” candidates) respectively. Specifically, we examined (1) the expression of immune checkpoint genes (ICGs), (2) the levels of immune cell fractions (ICFs), and (3) the ratios between immune cell fractions (ICFRs), as the predictive features for ICB response (**Fig. 5A**). More specifically, in the first category, we studied the expression of 79 ICGs in specific immune cell types, as reported in the literature (**Methods**). As a given ICG might be expressed in a few cell types, this resulted in a space of 219 gene-cell type pairs, out of which the expression state of 31 pairs in the TME could be predicted from the blood. In the second category, we focused on 9 ICFs in the blood whose levels were > 0 in the direct approach, and 6 ICFs in the TME whose levels could be predicted from the blood for the indirect, two step approach. In the third category, we comprehensively studied ICFRs in the form of (ICF_C1 ± ICF_C2) / (ICF_C3 + ICF_C4), where C1, C2, C3 and C4 are immune cell types. This space was composed of 3476 different ICFRs in the blood, out of which 55 ICFRs in the TME could be predicted from the blood. See **Methods** for more details and see **Supplementary material SM2** for a comprehensive list of items in each category.

**Figure 5.**
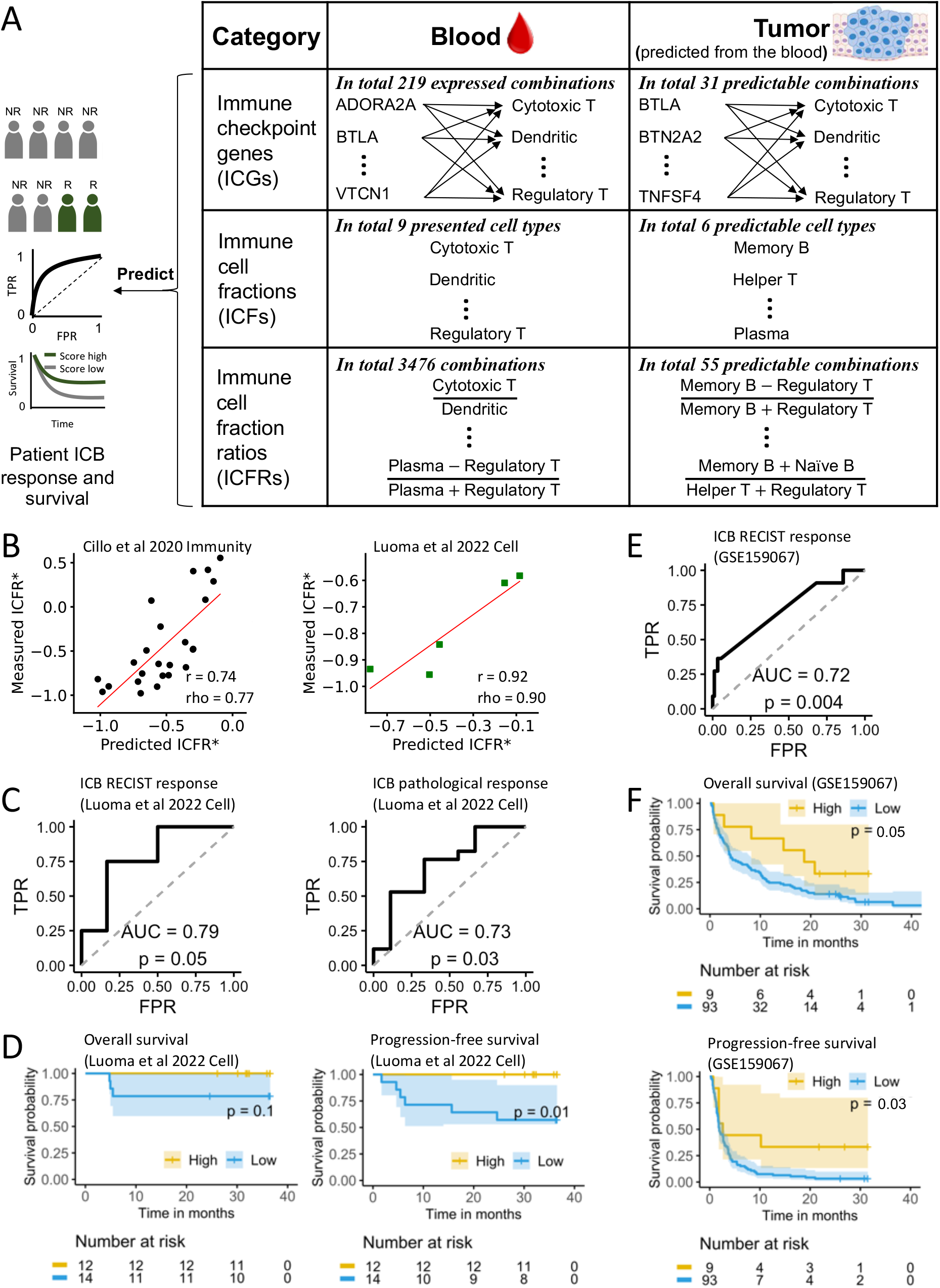
A new blood-predicted tumor immune signature, ICFR*, predicts both response and survival of HNSCC patients after ICB treatment. **A**. The predictive power for ICB response of three categories of immune information in the blood and in the TME are thoroughly evaluated, including the expression levels of key immune checkpoint genes in their corresponding cell-types, immune cell fractions, and the ratios between different immune cell fractions. **B-F**. Quantifying the prediction accuracy of the ICFR* signature (B_memory_ – T_reg_) / (B_memory_ + T_reg_). **B**. Correlation between the predicted ICFR* values and the measured ICFR* values in the TME in the matched blood/tumor training dataset (n = 25) and validation dataset (n = 5). **C**. ROC curve for predicting ICB RECIST response (n = 16) and pathological response (n = 26) by the TME ICFR* signature inferred from the blood scRNA-Seq. **D**. Overall survival and progression-free survival analyses of ICB-treated patients in ICFR*-high (score > quantile 50%) versus ICFR*-low (score ≤ quantile 50%) tumor groups for the PBMC scRNA-Seq dataset (GSE200996). **E**. ROC curve for predicting ICB response using the ICFR* computed from the deconvoluted bulk tumor RNA-Seq (GSE159067; n = 102). **F**. Overall and progression-free survival analysis of ICB-treated patients in ICFR*-high versus ICFR*-low tumor groups for the bulk tumor RNA-Seq dataset (GSE159067). Abbreviations: r, Pearson correlation coefficient; rho, Spearman correlation coefficient; TPR, true positive rate; FPR, false positive rate.

We next evaluated the correlation of these three categories of candidates and patients’ ICB response in the HNSCC dataset of [31]. Here the strength of the correlation was measured by the area under the receiver operating characteristic curve (AUC). The statistical significance of the correlation was calculated against the null hypothesis of “the AUC is equal to 0.5”. In category (1), we identified a few ICGs whose expression levels in the blood or in the TME were significantly associated with both ICB RECIST response (n=16) and ICB pathological response defined as “whether patients have 90% or less viable tumor after the ICB treatment” (n=26; **Methods**). The potential ICG signatures in the blood included LGALS9 expression in memory B cells, ICOSLG expression in naïve B cells, CD80 expression in monocytes, CD47 expression in regulatory T cells, and CD96 expression in memory B cells **(Supplementary Figure 8)**. The potential ICG signatures in the TME, whose expression levels were predicted from the blood, included LAG3 expression in NK cells, and ICOSLG expression in naïve B cells **(Supplementary Figure 9)**. In category (2), none of the ICFs, either from the blood or from the TME, was significantly associated with ICB response (**Supplementary material SM2**).

We hence focused on category (3), where there were three ICFR signatures in the TME that were quite highly predictive (**Supplementary material SM2**). The top predictive ICFR signature in the TME was (B_memory_ – T_reg_) / (B_memory_ + T_reg_), denoted as ICFR*. Firstly, ICFR* values in the TME could be well-predicted from the blood, with strong correlations between the predicted values and the measured values in both the training dataset of [29] (r = 0.75; rho = 0.77) and the validation dataset of [31] (r = 0.96; rho = 0.90; **Fig. 5B**). Second, inferred ICFR* values in the TME, which were predicted from the blood, were significantly associated with patients’ ICB response (positively correlated with ICB response; RECIST AUC = 0.79, p = 0.05; pathological AUC = 0.73, p = 0.03; **Fig. 5C**). Notably, as was the case with the exhausted T-cell signature, the ICFR* scores estimated directly from the blood were not significantly associated with ICB response (RECIST AUC = 0.54, p = 0.43; pathological AUC = 0.62, p = 0.17; **Supplementary Fig. 10**), in contrast to those inferred in the TME from the blood. Remarkably, none of the 3476 ICFRs in the blood was significantly associated with ICB response (**Supplementary material SM2**), which further underscored the importance of predicting ICB response via the two-step approach. Finally, while signatures from category (1) were not significantly associated with patients’ survival after ICB treatment **(Supplementary Figures 8, 9)**, which might be due to the short follow-up time issue of this dataset as mentioned above, the ICFR* scores in the TME were predictive of patients’ survival after ICB treatment (OS, p = 0.1; PFS, p = 0.01; **Fig. 5D**).

To further test the robustness of the ICFR* signature in predicting ICB response, we analyzed an additional large cohort. This cohort was a recently released dataset [35], consisting of bulk tumor RNA-Seq of 102 advance HNSCC patients treated with ICB. Though it did not include matched single cell data, we were still able to use it to test the predictive power of the ICFR* signature, as described below. To calculate the ICFR* values in the TME in each of these tumor samples, we inferred the relative abundance of each immune cell type by deconvolving the bulk tumor RNA-Seq data via CIBERSORT [36]. Remarkably, the ICFR* scores were also predictive of ICB response in this independent dataset (AUC = 0.72, p = 0.004; **Fig. 5E**), which was much higher than the predictive power obtained using the FDA-approved TMB biomarker in HNSCC (AUC=0.56, p =0.74; n = 69; **Supplementary Figure 11**; data from the MSK-IMPACT cohort [37]). Additionally, ICFR* values significantly predicted patients’ survival after ICB treatment (OS, p = 0.05; PFS, p = 0.03; **Fig. 5F**). It was also worth noting that ICFR* didn’t not predict HNSCC patients’ survival in TCGA, where patients were not treated with ICB (**Supplementary Figure 12**), demonstrating its specificity for predicting ICB response.

For completeness, we also evaluated the predictive power of the two other ICFR signatures that ranked high on our list. They were (B_memory_ – T_reg_) / (T_reg_ + Plasma) (positively correlated with ICB response; RECIST AUC = 0.79, p = 0.05; pathological AUC = 0.73, p = 0.03), and B_memory_ / (T_reg_ + Plasma) (positively correlated with ICB response; RECIST AUC = 0.79, p = 0.05; pathological AUC = 0.73, p = 0.03) (**Supplementary Fig. 13**). Notably, these two ICFR signatures were also able to significantly predict the ICB response in the bulk dataset (AUC = 0.72, p = 0.004 and AUC = 0.63, p = 0.007, respectively), but were less predictive of patients’ survival after ICB treatment (**Supplementary Fig. 13**).

## Discussion

Cancer is a systemic disease that affects the entire body, altering the composition and function of the immune system as a whole [38]. Charting the immune status of the TME in individuals is crucial for understanding cancer progression, drug response and resistance, which can help promote precision medicine [39]. Due to the challenges and risks associated with tumor tissue biopsies, characterizing the TME from liquid biopsies has become one of the most important directions in cancer prognosis, stratification, and monitoring [40]. Leveraging the advantage of single cell sequencing techniques, here, we have established a machine learning framework utilizing the global expression profiles of major immune cells in the peripheral blood, derived from the CD45+ scRNA-Seq data, to infer the immune status of the TME in HNSCC patients (**Fig. 1D**). To the best of our knowledge, the immune status of the TME has not yet been comprehensively predicted from the blood previously.

Our results testify that ICFs of major immune cell types in the TME of HNSCC primary tumors can be inferred from the blood. The most predictable three immune cell types are memory B cells, regulatory T cells and helper T cells on the independent validation data (**Fig. 2C**). Coincidently, all these cell types have been shown to play important roles in regulating cancer prognosis and drug resistance in HNSCC patients [41, 42]. Additionally, we show that the expression levels of approximately 20 – 50% of the genes expressed in different immune cell types in the TME is strongly correlated between the blood and the TME in (mostly) the same cell-types and can be predicted from the blood. Furthermore, our findings suggest that the clinical information significantly contributes to the predictability of both ICFs and gene expression levels of immune cells in the TME. Interestingly, different clinical variables are found to contribute to the prediction of the expression profiles of different immune cells. For example, among all 5 clinical variables, HPV infection status is found to predict the expression of the largest number of genes in dendritic cells and all T cells; while tobacco use information is the largest contributor for innate immune cells (i.e., monocytes, macrophages, NK cells), and alcohol use is the largest contributor for B cells and plasma cells (**Fig. 3B**). This suggests that different clinical variables may have different impacts on the gene expression levels of different cell types.

Turning to study the ability to predict patients’ response to ICB from the inferred ICFs and gene expression in the TME, we show that one of the most well-known pan-cancer immune signatures for immunotherapy, the exhausted T-cell signature, can be predicted from the blood single-cell transcriptomics in HNSCC and the inferred exhausted T-cell signature from the blood can predict patients’ ICB response (**Fig. 4A, B**). Moreover, we identify a new immune signature, ICFR*, whose inferred TME scores from the blood are more strongly associated with patients response. Specifically, the ICFR* predicts both ICB response and survival on both single-cell and bulk datasets (**Fig. 5B-F**).

Additionally, it is worth noting that the predictive power of ICFR* holds across different patient populations: the single-cell cohort contains only newly diagnosed oral cavity squamous cell carcinoma patients, who have received the ICB as a neoadjuvant therapy before the surgery; whereas, in the bulk cohort, the ICB treatment has been used as a main therapy for patients with advanced HNSCC, and more than half of the patients have received at least one line therapy before the ICB treatment. In addition, ICFR* is not predictive of the survival of HNSCC patients that are not treated with ICB therapy, in the TCGA cohort. Taken together, these results demonstrate that the prediction ability of ICFR* is fairly robust and also specific to immunotherapy treatment. It is also worth noting that in contrast to the predictive power of inferred ICFR* scores in the TME, none of the thousands of possible ICFR scores in the blood are predictive of ICB response, which highlights the importance of constructing a two-step predictor to predict ICB treatment outcome, i.e., first learning about the immune status of the TME from the blood data, and then using the latter to further predict the treatment outcome. Notably, we find that B cells play a crucial role in the three top ICFR signatures identified in this research, which is in line with recent studies that demonstrate the importance of B cells on ICB treatment outcome [43-45].

The present study has some limitations that should be acknowledged. Firstly, the sample size used to train the models is relatively small, as the study relies on yet costly scRNA-Seq data. Currently, there is no larger publicly available dataset that contains matched tumor-PBMC scRNA-Seq data. Although by constructing very compact predictors, the risk of overfitting has been reduced and the resulting models have shown reasonable performance on an independent validation dataset, larger cohorts are still needed to further test and validate the results. Specifically, as the predictive power of inferred TME ICFR* signature has only been tested in one single-cell dataset, more PBMC cohorts are needed to further test and validate this potentially clinically relevant biomarker. Secondly, the focus of this study is on the relative fractions of immune cells, as the fractions of different immune cells out of all cells (including malignant cells and other hematopoietic cells) are not available in current datasets. Such future studies may potentially further enhance ICB response prediction. Thirdly, we observe an interesting correlation pattern between ICFs in the TME and blood (**Fig. 2A**), which suggests that there may be possible interactions between immune cells in the TME and blood that may have been previously overlooked. To predict ICFs for a specific immune cell type in the TME, we used the most highly correlated cell type in the blood to build our predictive models as a starting point. This can be conceived as a prior feature selection step that may introduce a potential bias when evaluating the model in cross validation. However, given the limited sample size, the current strategy is the best option one could adopt. If we would have instead tested the ICFs of all cell types and all the clinical variables, the number of candidate models would have greatly exceeded the training sample size, making that analysis implausible. In the future, as more data becomes available, it will be important to revisit and investigate the robustness and underlying mechanisms of the relationship between ICFs in the TME and blood in HNSCC. With future increased sample size, one could then also use more robust cross-validation models to test and validate the prediction results in more depth.

Transcriptome-based biomarkers are showing increasing power in predicting cancer prognosis and immunotherapy response [43, 44]. Here we have shown the feasibility of constructing predictive models of immune cell fractions and gene expression levels in the TME based on matching blood single-cell transcriptomics. We further show that the latter TME blood-derived information is in turn predictive of patients’ ICB response. These results offer a new and promising way to extend the realm of liquid biopsies beyond ctDNAs and CTCs to that of blood SC transcriptomics. Taken together, our work represents a promising initial step towards utilizing single-cell transcriptomics based liquid biopsies to accurately predict the immune status of patients’ tumors from their blood, facilitating personalized cancer precision therapy in the future.

## Materials and methods

### Data collection

In this study, the primary sources of the data are obtained from NCBI Gene Expression Omnibus (GEO; http://www.ncbi.nlm.nih.gov/geo/). Additional information regarding the datasets used in this study can be found in **Table S1**.

### Single-cell RNA-seq data analysis

Seurat (v4.0.1, R package) [46] was used for data processing and cell clustering. Cells with insufficient number of genes (< 200) or greater than 20% of UMIs mapped to mitochondrial genes were removed. Data integration across different samples was accomplished with Harmony [47]. The *‘NormalizeData’* function with parameters: *normalization.method = “LogNormalize”* and *scale.factor = 10000* was applied to normalize the expression levels of genes in each single cell. The *‘FindVariableFeatures’* function with the *‘vst’* method was utilized to identify 2,000 highly variable genes. The *‘ScaleData’* function with default parameters was used to scale and center gene expression matrices. To perform clustering, principal component analysis (PCA) dimensionality reduction was first conducted with *‘RunPCA’* function and the first 20 principal components were selected to construct the shared nearest neighbor (SNN) graph with *‘FindNeighbors’* function, and then clusters were determined using the Louvain algorithm with *‘FindClusters’* function. CellTypist [30] with the default model (*Immune_All_Low.pkl*) and majority voting refinement was used to do automatic cell type annotation with the known cell type labels to predict the identities of cell clusters. Then, we manually checked whether the annotations are reliable by examining the top ranked differentially expressed genes of each cluster which were obtained with *‘FindAllMarkers’* function with default parameters but with set *min.pct = 0.25*. The uniform manifold approximation and projection (UMAP) was finally applied to visualize the single cell transcriptional profile in 2D space.

### Differential cell abundance and gene expression analysis

Differential cell abundance analysis was conducted with edgeR [48] to identify which cell types were differentially enriched in tumor tissue or in the blood. ‘*estimateDisp()*’ function was used to estimate the negative binomial dispersion for each cluster. ‘*glmQLFit()’* function was used to estimate the quasi-likelihood dispersion. *‘glmQLFTest()’* function was used to test for differences in abundance between tumor tissue and blood with FDR < 0.05.

In this study, the gene expression of a specific immune cell type was defined as “pseudo-bulk” expression profiles generated by summing counts together for all individual cells of the same cell type in a sample. “Pseudo-bulk” samples with insufficient cells (< 10) were removed. As a result, 10 cell types (cycling T cells, cytotoxic T cells, dendritic cells, helper T cells, memory B cells, monocytes, naive B cells, NK cells, plasma cells, regulatory T cells) were retained for differential gene expression analysis. ‘*pseudoBulkDGE()*’ function from scran (v1.22.1, R package) was used to perform the differential gene expression analysis to find significantly up and downregulated genes with FDR < 0.05 and log2 fold change (FC) = 0. This approach generated for each cell type a list of genes over and under-expressed in the tumor compared to the peripheral blood. GO enrichment analyses of the resulting gene lists were performed with R package *clusterProfiler* (v 4.2.2).

### Machine learning for prediction of the ICFs in the TME

In order to predict the ICFs in the TME from the blood, for each immune cell type in the TME, we first identified the most predictive cell type in the blood using the highest Pearson correlation coefficient (r_max_) of ICFs with the target cell types in the TME. If the correlation between a cell type in the TME with the identical cell type in the blood was higher than r_max_ * 0.9, then the identical cell type was used as the most predictive cell type. To reduce the impact of potential sampling bias, the correlation was calculated using 1000-replicate bootstrapping.

Then, for each cell type in the TME, we constructed a 1-variable linear regression baseline model by using the ICF of the corresponding most predictive cell type in the blood (as the only variable; Model 1). Additionally, we constructed five 2-variable linear regression models by using the ICF of the corresponding most predictive cell type in the blood (as the first variable) and one of the five clinical variables (as the second variable; Models 2 - 6).

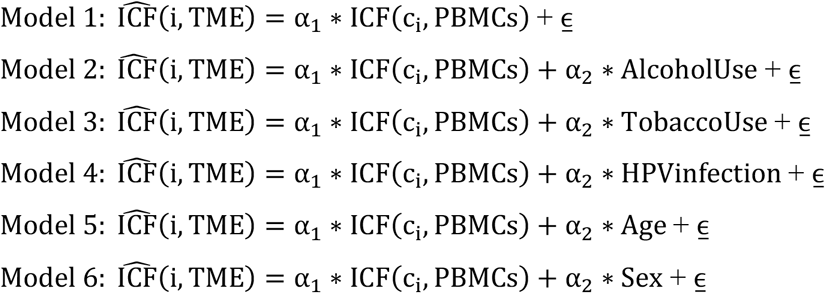

where i is the i-th immune cell type in the TME; c_i_ is the corresponding most predictive cell type in the blood.

To reduce the risk of overfitting, elastic-net regularization was used during model training. Finally, for each cell type, among all 6 models we selected the model with the best performance as the final model.

To perform cross validation, the data were randomly split (80%: 20%) to do the training and testing for 1000 replicates, and results were computed and analyzed over all 1000 replicates.

The ICF of a specific immune cell type in the TME is defined predictable when the mean Pearson correlation coefficients between the predicted ICFs and the true ICFs on the 20% test sets are greater than 0.3 (r > 0.3), mean normalized mean absolute error less than 1 (NMAE < 1) with a Benjamini-Hochberg adjusted p-value < 0.05 (FDR < 0.05) over 1000 replicates. More details are in the following section of “*Statistical test to identify predictable ICFs and genes*”.

For model evaluation on the validation dataset and model application in predicting patient ICB response, the mean values of the parameters (i.e., coefficients and intercepts of the linear models) of the final models trained over 1000 replicates were used.

### Machine learning for prediction of the gene expression levels in immune cells in the TME

Similar to the prediction of ICFs, we used 1-variable and 2-variable linear regression models to predict expression levels of expressed genes in different immune cell types in the TME. In this case, “expressed genes” are defined as genes that have non-zero expression values in more than half of the samples.

Specifically, for each immune cell type in the TME, we first identified the most predictive cell type in the blood based on the Pearson correlation coefficient of the overall gene expression levels across samples. Then, we used six linear regression models to predict the expression level of a specific gene in a specific immune cell type in the TME.

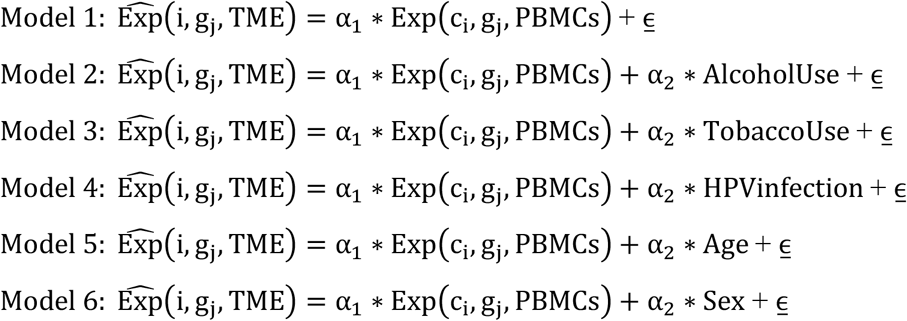

where i is the i-th immune cell type in tumor tissue; g_j_ is the j-th expressed gene; c_i_ is the corresponding most predictive cell type in PBMCs identified by correlation of overall gene expression levels with i.

Elastic-net regularization was used during model training. Finally, for each gene, among all the 6 models we selected the model with the best performance as the final model. The results were computed, evaluated, and analyzed over 1000 replicates.

### Statistical test to identify predictable ICFs and genes

Pearson correlation coefficients between the predicted and true values of the ICF/gene expression in the TME on the test data over 1000 replicates were calculated for each cell type/gene, and were tested against the null hypothesis of:

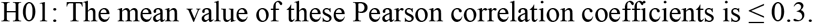

Similarly, the normalized mean absolute errors between the predicted and true values over all 1000 replicates were tested against the null hypothesis of:

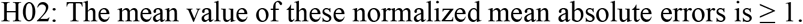

We applied FDR ≤ 0.05 to both hypotheses (H01 and H02) to select the predictable ICFs and genes.

Regarding identifying the clinical variables that have the highest and significant contribution to increasing the predictive power of each model, we compared results from models 2 - 6 with result from model 1 using the Mood’s median test.

### Functional enrichment analysis

R package clusterProfiler (v 4.2.2) was used to perform functional enrichment analysis of predictable gene lists including enrichment analysis of GO and enrichment analysis of KEGG (Kyoto Encyclopedia of Genes and Genomes), as well as enrichment analysis of hallmarks gene sets and reactome subset of canonical pathways downloaded from MSigDB (http://www.gsea-msigdb.org/).

### Identification of blood-predictive signatures for prediction of ICB response

To identify potential immune signatures that can be learned from the blood and are predictive of HNSCC patients’ ICB response, we comprehensively examined three categories of candidate immune information in the blood and in the TME respectively, i.e., ICGs, ICFs, and ICFRs (see **Supplementary material SM2** for comprehensive lists of them). Note that all the three categories of candidate immune information in the TME are predicted from the blood scRNA-Seq data using the machine learning models described above.

Regarding ICGs, in total 79 ICGs were collected from literature review [9, 49-52]. We first checked the expression levels of them in different immune cell types in the blood. There are 219 expressed pairs in the blood, among which 31 pairs in the TME can be predicted from the blood (with criteria of r > 0.3 and FDR < 0.05 on the training dataset; r> 0.3 on the validation dataset). Regarding ICFs, 9 are present in the blood (ICF > 0) and 6 in the TME can be predicted from the blood (with criteria of r > 0.3 and FDR < 0.05 on the training dataset; r> 0.3 on the validation dataset; **Fig. 2C**). Regarding ICFRs, there are in total 3476 different combinations in the form of (ICF_C1 ± ICF_C2) / (ICF_C3 + ICF_C4) in the blood, where C1, C2, C3, and C4 are immune cell types, and 55 in the TME are significantly predictable (with criteria of r > 0.3, P ≤ 0.05; rho > 0.3, P ≤ 0.05 in both the training dataset and the validation dataset). As C1, C2, C3, and C4 are not necessarily different from each other, the form of (ICF_C1 ± ICF_C2) / (ICF_C3 + ICF_C4) can actually include all the following forms: ICF_C1 / ICF_C2, ICF_C1 / (ICF_C2 + ICF_C3), (ICF_C1 ± ICF_C2) / ICF_C3, and (ICF_C1 ± ICF_C2) / (ICF_C3 + ICF_C4).

We then ranked these candidate predictors in each category based on their correlation with ICB response using data from [31]. Here the strength of the correlation was measured by the area under the receiver operating characteristic curve (AUC). The Mann-Whitney U test was used to calculate the p-values against the null hypothesis of “the AUC is equal to 0.5” [53]. AUCs and corresponding p-values for predicting both the ICB RECIST response (n=16) and the ICB pathological response (n=26) were calculated. Predictors with p ≤ 0.05 for both RECIST response and pathological response were considered as potential signatures.

### Survival analyses

All Kaplan–Meier analyses were performed by comparing the survival of patients with high signature scores (> median) to those with low scores (≤ median) using a two-sided log-rank test.

## Code availability

All codes that are necessary to reproduce all the results in the paper are implemented in Python and R and are publicly available in GitHub: https://github.com/yingstat/TIMEP.

## Acknowledgements

This work utilized the computational resources of the NIH HPC Biowulf cluster (http://hpc.nih.gov).

## Supplementary information

**Supplementary Table 1.** Datasets used in this study.

| Data source | Samples | Description | Purpose | Reference |
| --- | --- | --- | --- | --- |
| GSE139324 | 26 | Matched tumor-PBMC CD45+ scRNA-Seq | Model training | [29] |
| GSE200996 | 5 | Matched tumor-PBMC CD45+ scRNA-Seq | Model validation | [31] |
|  | 27 | PBMC CD45+ scRNA-Seq, patients' ICB response | Signature identification | [31] |
| GSE159067 | 102 | Bulk tumor RNA-Seq, patients' overall survival and progression-free survival, ICB response | Signature validation | [35] |
| MSK-IMPACT | 69 | TMB, patients' ICB response | Predictive power comparison | [37] |
| TCGA | 520 | Bulk tumor RNA-Seq, patients' overall survival and progression-free survival | Survival analysis | [54] |

**Supplementary Table 2.** Functional enrichment of predictable genes in the 11 immune cell types in the TME. The counts represent the number of immune cell types in which the corresponding significantly enriched term is present. The 15 most frequently present GO, KEGG, HALLMARK and REACTOME terms (all with FDR < 0.05) are shown, respectively.

| GO term | Counts | KEGG term | Counts | HALLMARK term | Counts | REACTOME term | Counts |
| --- | --- | --- | --- | --- | --- | --- | --- |
| Adaptive immune response based on somatic recombination of immune receptors built from immunoglobulin superfamily domains | 11 | Cellular senescence | 11 | TNFA signaling via NFkB | 11 | Class_I MHC mediated antigen processing presentation | 11 |
| Aromatic compound catabolic process | 11 | Epstein-Barr virus infection | 11 | Allograft rejection | 10 | Neutrophil degranulation | 11 |
| Cellular nitrogen compound catabolic process | 11 | Graft-versus-host disease | 11 | MYC targets v1 | 10 | Transcriptional regulation by TP53 | 11 |
| Heterocycle catabolic process | 11 | Human T-cell leukemia virus 1 infection | 11 | Interferon gamma response | 9 | Activation of the mRNA upon binding of the cap binding complex and EIFS and subsequent binding to 43s | 10 |
| Leukocyte cell-cell adhesion | 11 | Allograft rejection | 10 | Interferon alpha response | 8 | Cellular response to starvation | 10 |
| Organic cyclic compound catabolic process | 11 | Antigen processing and presentation | 10 | Oxidative phosphorylation | 8 | Eukaryotic translation elongation | 10 |
| Positive regulation of T cell activation | 11 | Coronavirus disease - COVID-19 | 10 | DNA repair | 7 | Eukaryotic translation initiation | 10 |
| Protein folding | 11 | Endocytosis | 10 | MTORC1 signaling | 7 | HIV infection | 10 |
| Response to endoplasmic reticulum stress | 11 | Kaposi sarcoma-associated herpesvirus infection | 10 | Unfolded protein response | 7 | Influenza infection | 10 |
| Response to virus | 11 | Phagosome | 10 | E2F targets | 6 | Interferon signaling | 10 |
| Ribosome assembly | 11 | Ribosome | 10 | G2M checkpoint | 4 | Metabolism of amino acids and derivatives | 10 |
| T cell activation | 11 | Salmonella infection | 10 | Mitotic spindle | 4 | mRNA splicing | 10 |
| Cytoplasmic translation | 10 | Spliceosome | 10 | Apoptosis | 3 | Nonsense mediated decay NMD | 10 |
| Establishment of protein localization to organelle | 10 | Th17 cell differentiation | 10 | Complement | 3 | Processing of capped intron containing pre mRNA | 10 |
| Histone modification | 10 | Type I diabetes mellitus | 10 | IL6 JAK STAT3 signaling | 3 | Regulation of expression of SLITS and ROBOS | 10 |

**Supplementary Figure 1.**
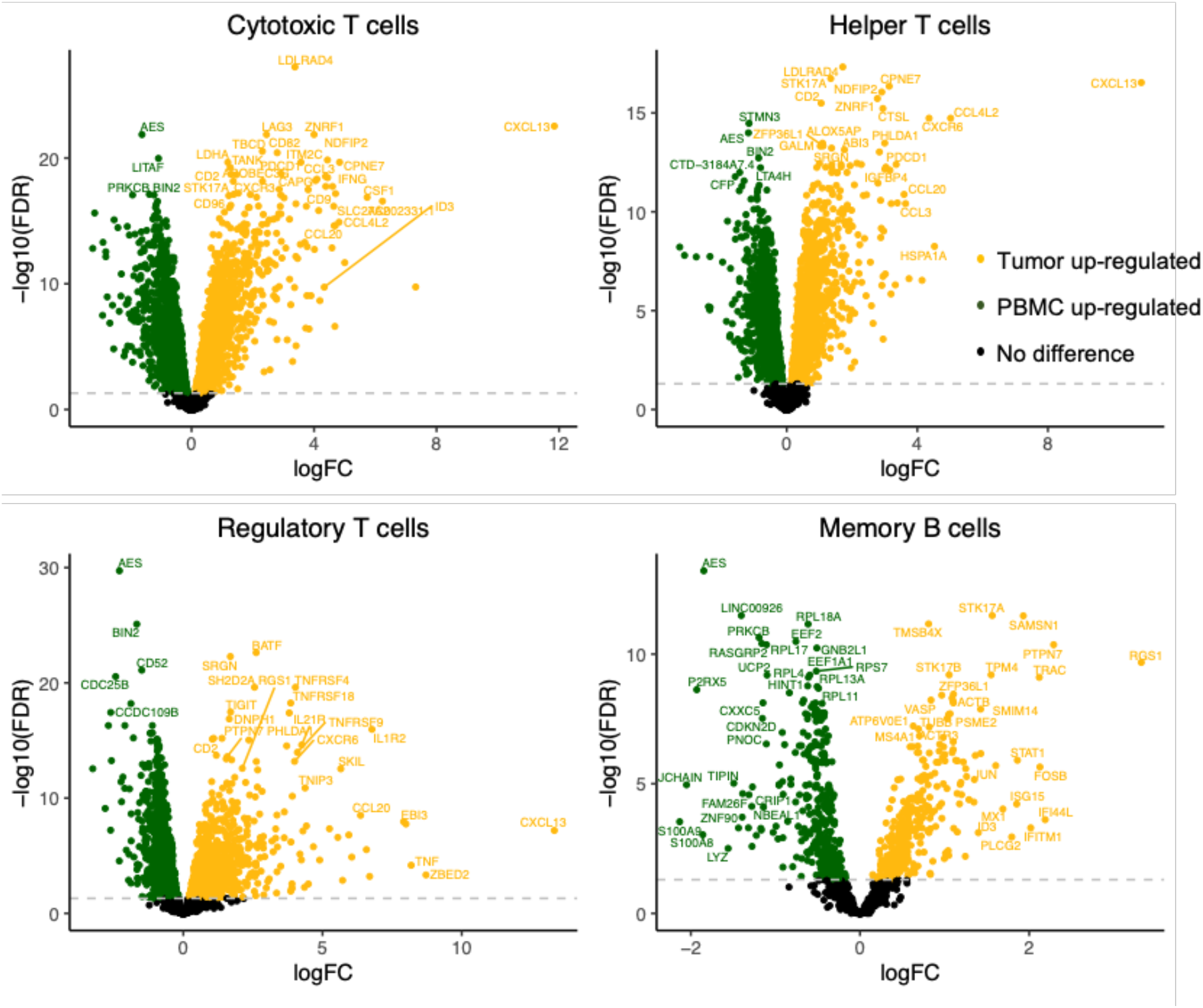
Immune response related genes are more highly expressed in the TME compared to the blood.

**Supplementary Figure 2.**
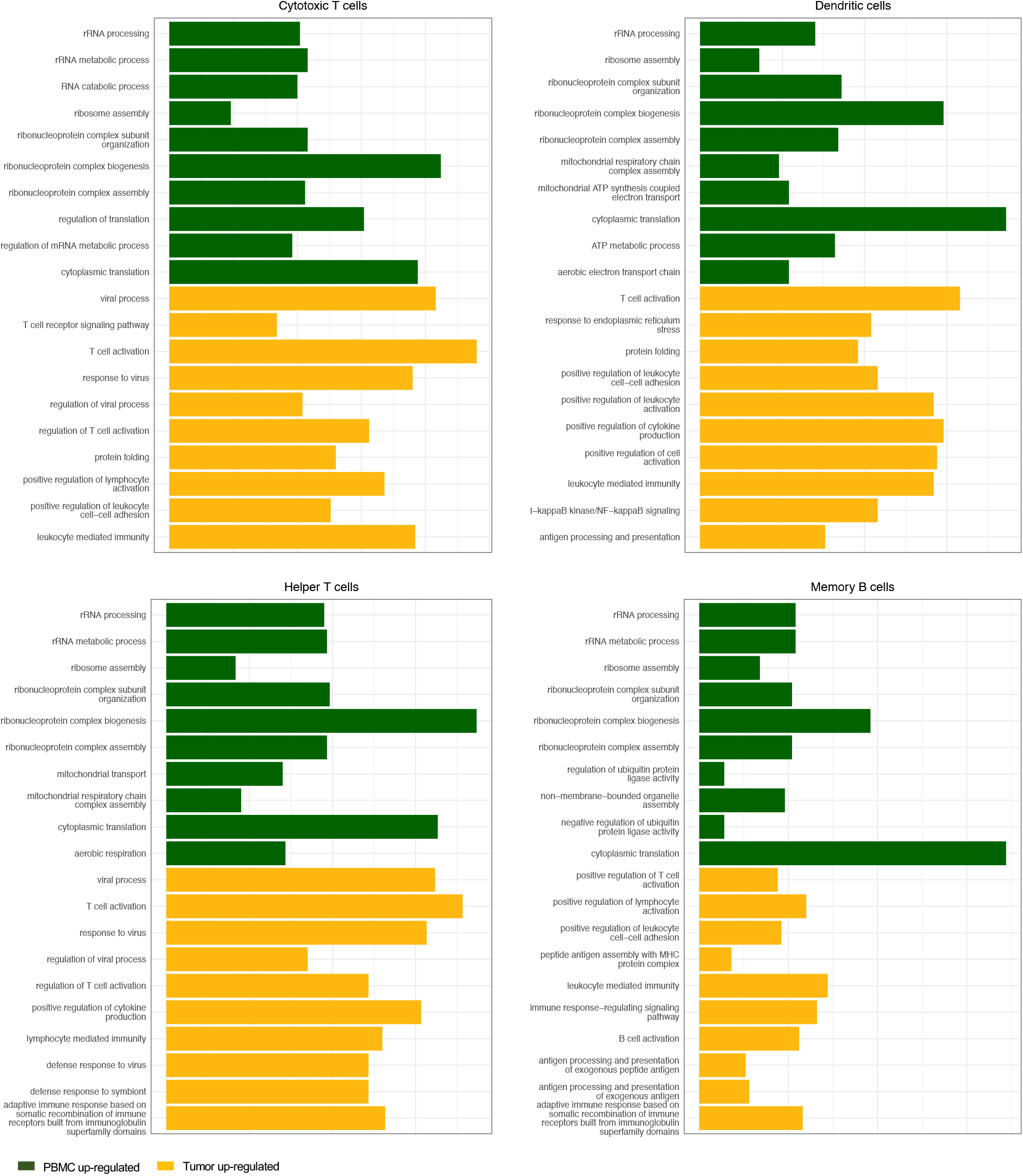
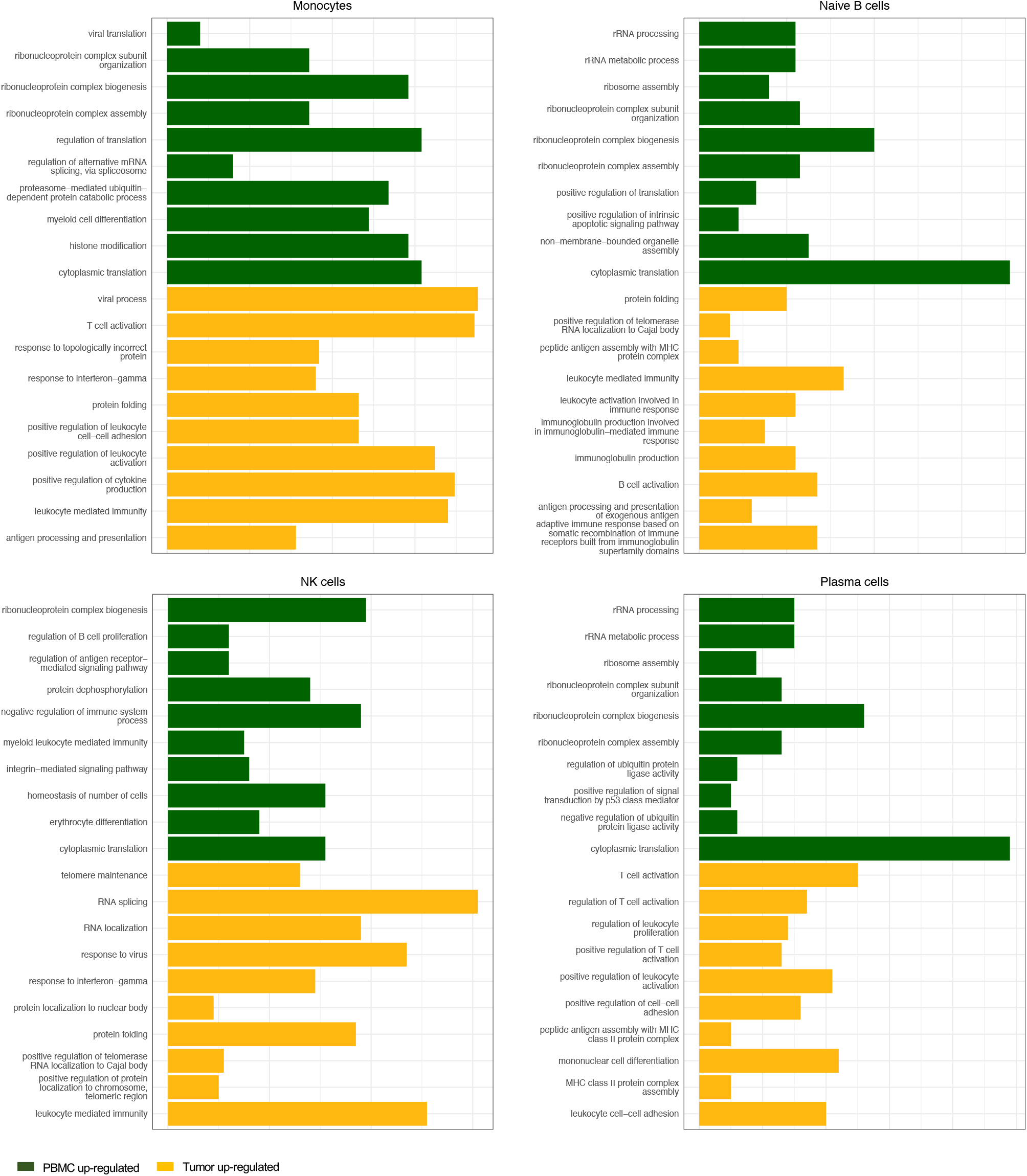
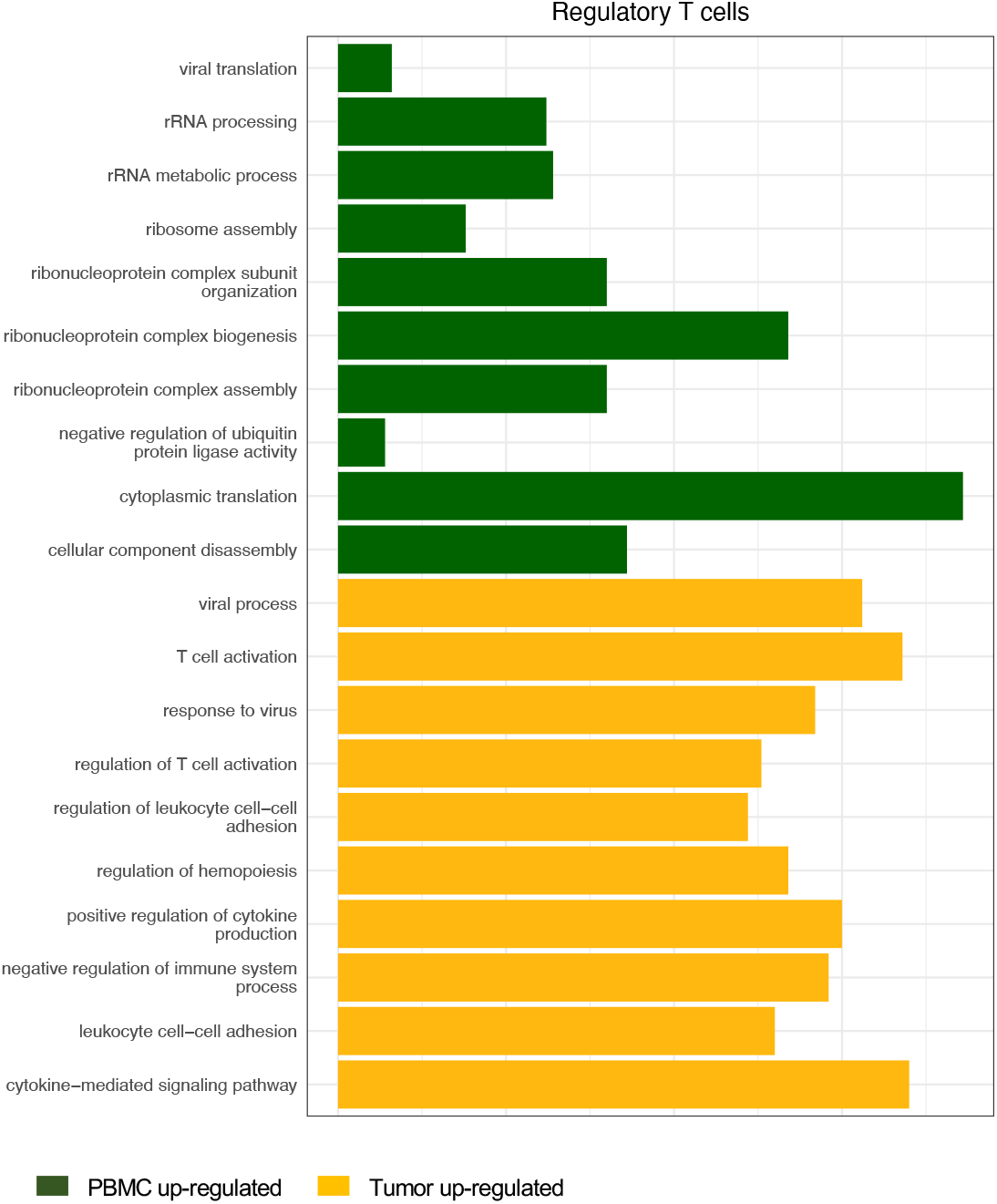
Functional enrichment of up-regulated genes in different immune cells in the TME and in the blood.

**Supplementary Figure 3.**
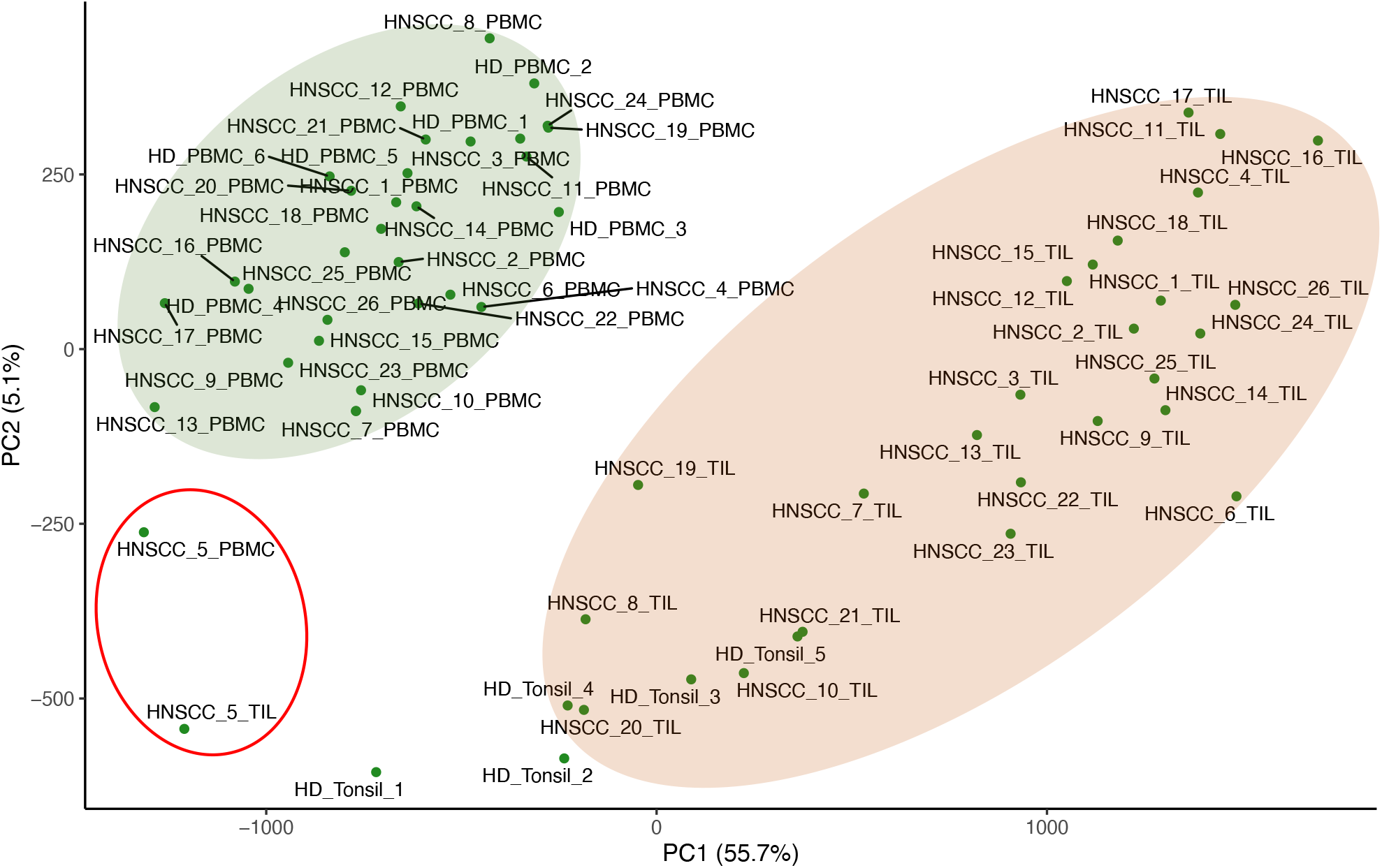
A PCA plot of the pseudo-bulk gene expression data from CD45+ PBMC and primary tumor tissue samples from 26 HNSCC patients and 11 healthy donors. Sample 5 from the HNSCC patients is excluded from further analysis as an outlier, which means that it is significantly different from the other samples and may not be representative of the group. The data is from [29].

**Supplementary Figure 4.**
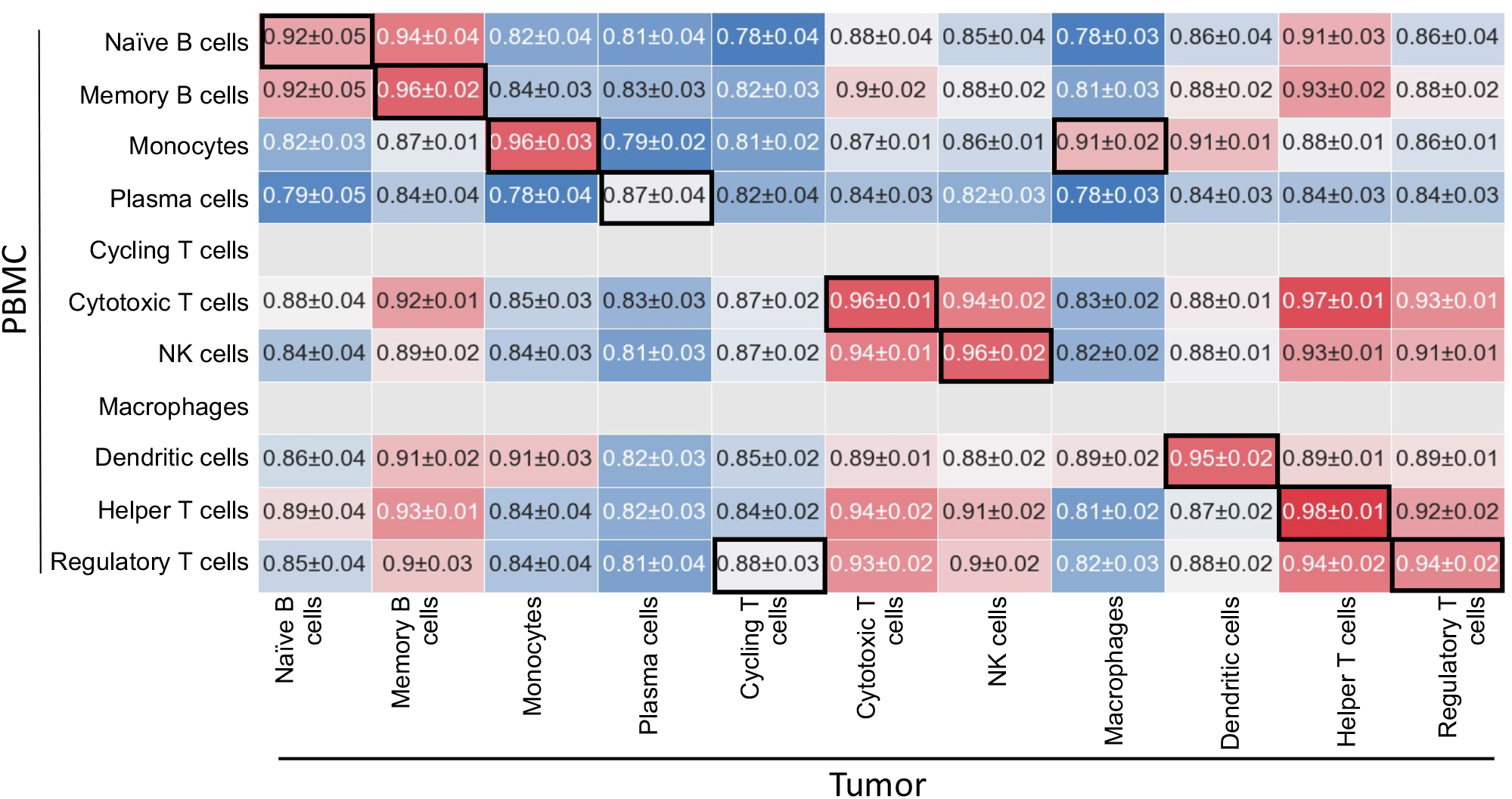
Correlation matrix that compares whole-genome gene expression levels of major immune cell types in the TME to those in the blood. The cells in the TME are listed as columns and the cells in the blood are listed as rows. The matrix shows the correlation between the overall gene expression levels of each cell type in the TME and the corresponding cell type in the blood. The cell type in the blood that best correlates with a given cell type in the TME is marked in a black box. The correlation is calculated by flattening the expression levels of all genes in a cell type (mean ± s.d., where s.d. are calculated across the 25 samples).

**Supplementary Figure 5.**
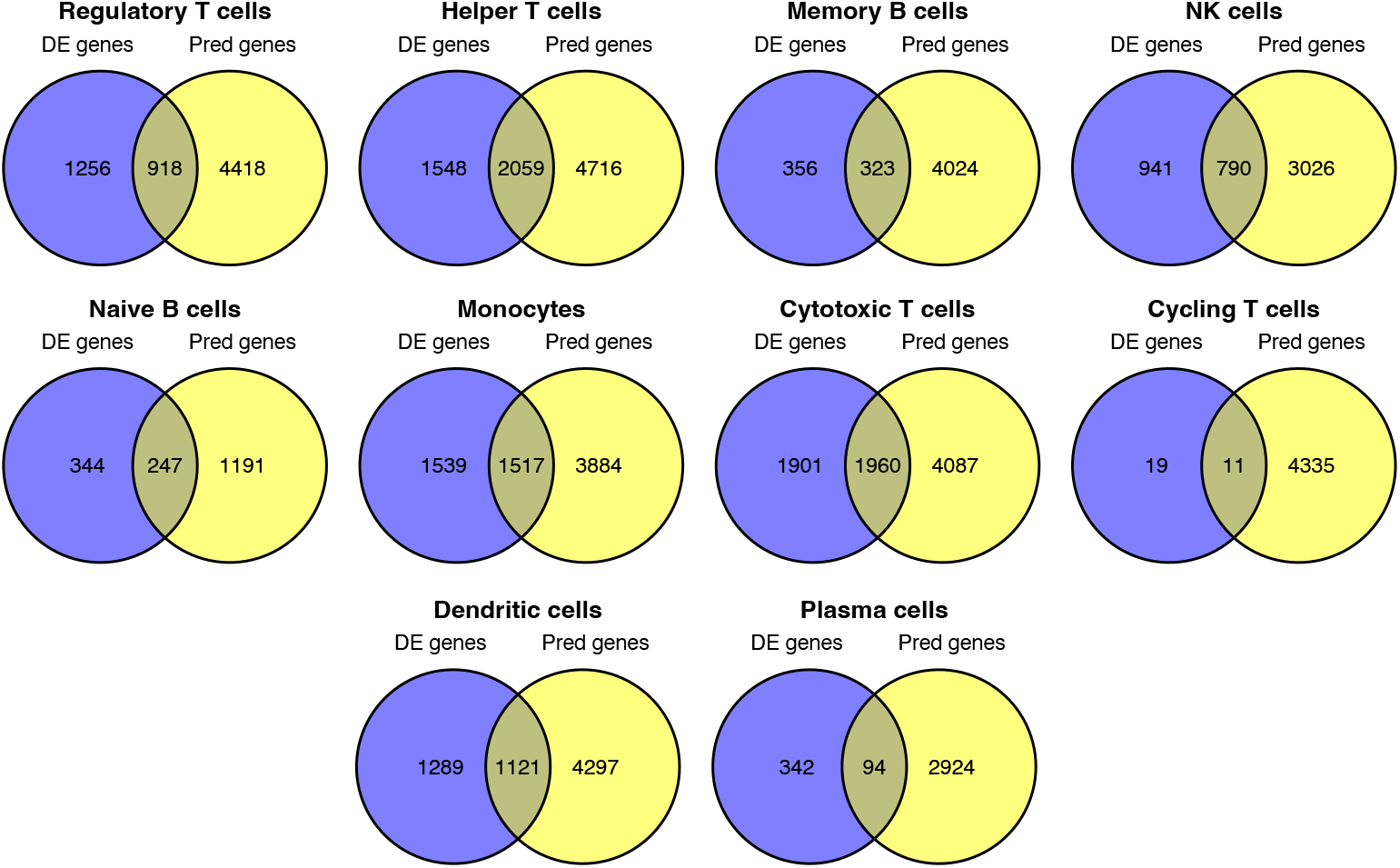
The expression levels of 22-57% differentially expressed genes between immune cells in the TME and in the blood can be predicted from the blood. Note: DE genes, differentially expressed genes in immune cells between the TME and the blood; Pred genes, genes whose expression levels in immune cells in the TME can be predicted from the blood.

**Supplementary Figure 6.**
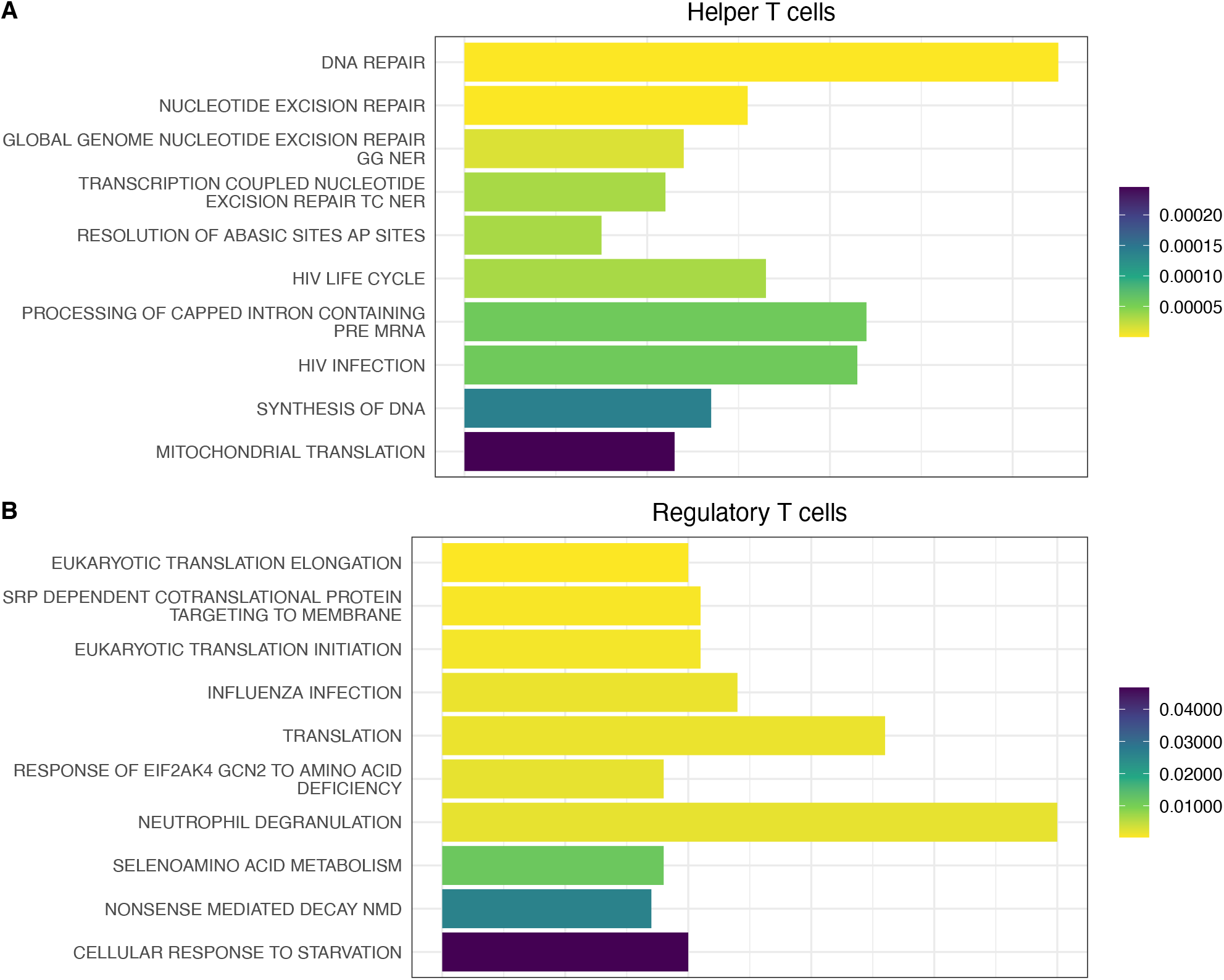
Reactome functional enrichment of genes in helper T cells (A) and regulatory T cells (B) that are most accurately predicted based on the HPV infection status information.

**Supplementary Figure 7.**
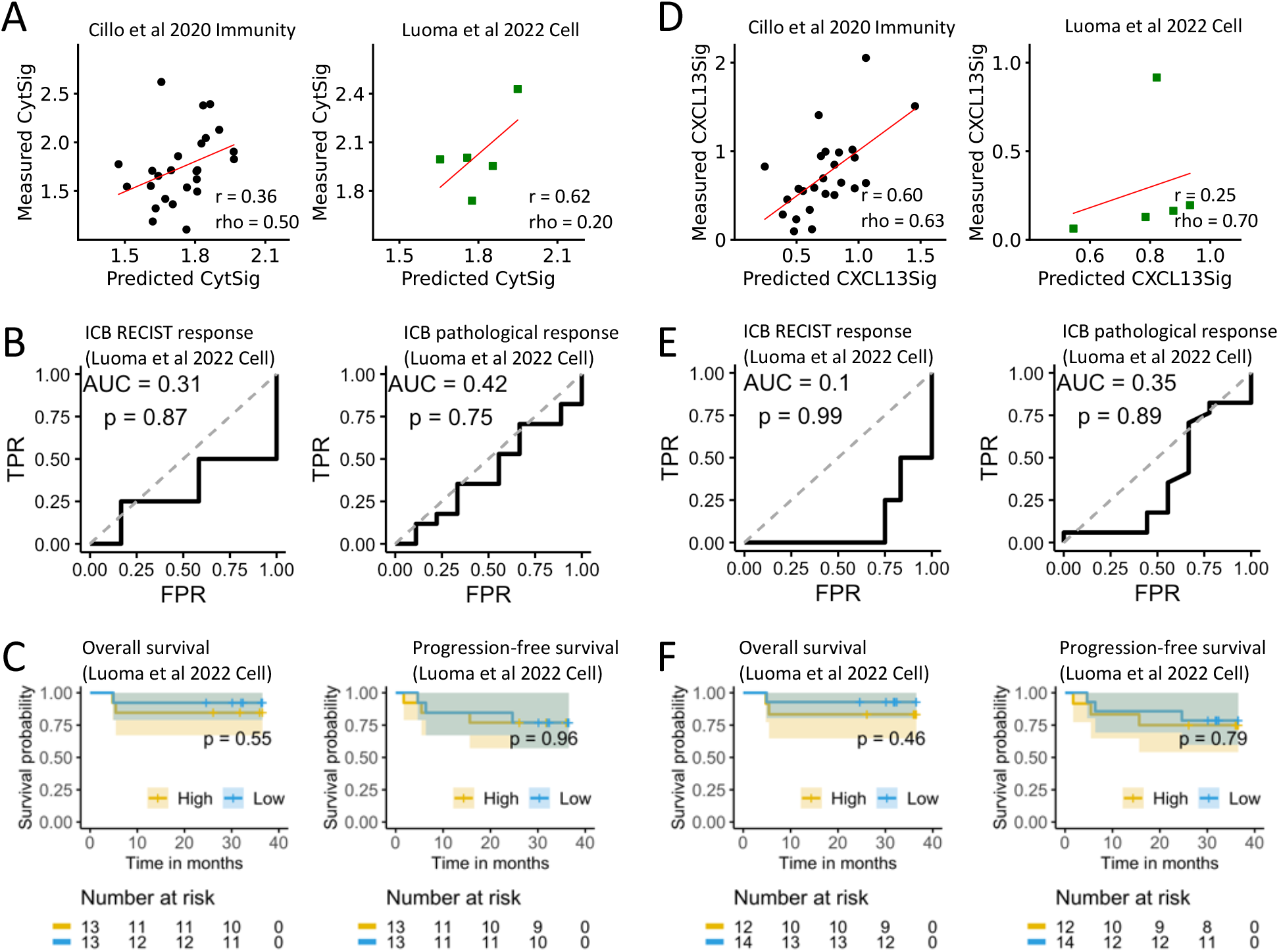
The cytolytic T-cell and the CXCL13 signatures in the TME are less predictable from the blood. **A**. Correlation between the predicted and the measured cytolytic T-cell signature scores in TME in the matched blood/tumor training dataset (n = 25) and validation dataset (n = 5). **B**. ROC curve for predicting ICB RECIST response (n = 16) and pathological response (n = 26) by the predicted cytolytic T-cell signature scores. **C**. Overall survival and progression-free survival analyses of ICB-treated patients in cytolytic-high (cytolytic score > quantile 50%) versus cytolytic-low (cytolytic score ≤ quantile 50%) tumor groups using the predicted cytolytic T-cell signature scores. Panels **D, E, F** are the same to A, B, C, respectively, except that the cytolytic T-cell signature is replaced by the CXCL13 signature. Abbreviations: r, Pearson correlation coefficient; rho, Spearman correlation coefficient; TPR, true positive rate; FPR, false positive rate.

**Supplementary Figure 8.**
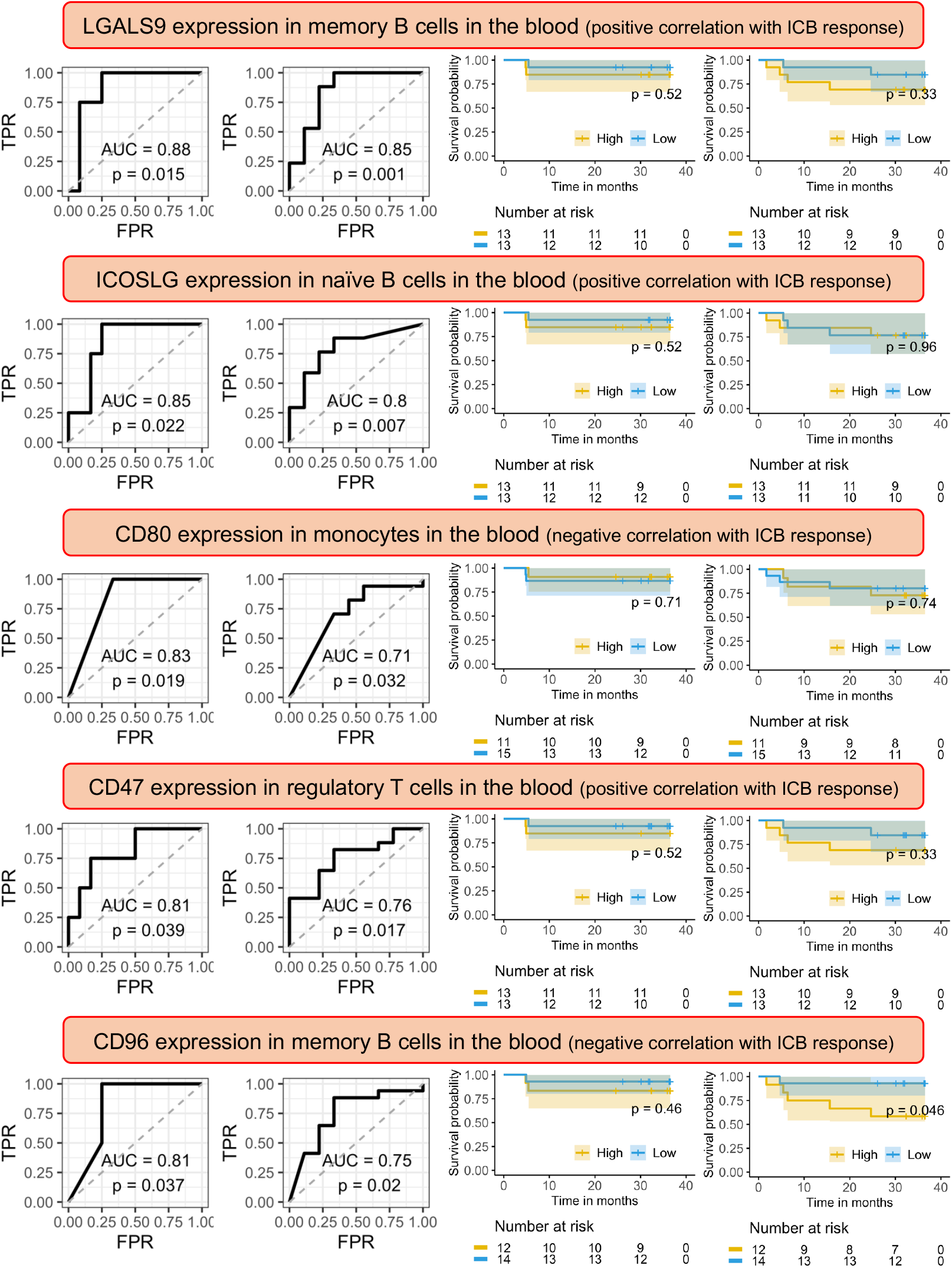
Five blood ICG signatures are found to predict ICB response but not constantly predict patient survival after ICB treatment in HNSCC patients. The names of the five signatures are shown in the figure. The first two columns are ROC curves of ICB RECIST response and pathological response respectively. The last two columns are overall survival and progression-free survival after ICB treatment respectively. The data is from [31].

**Supplementary Figure 9.**
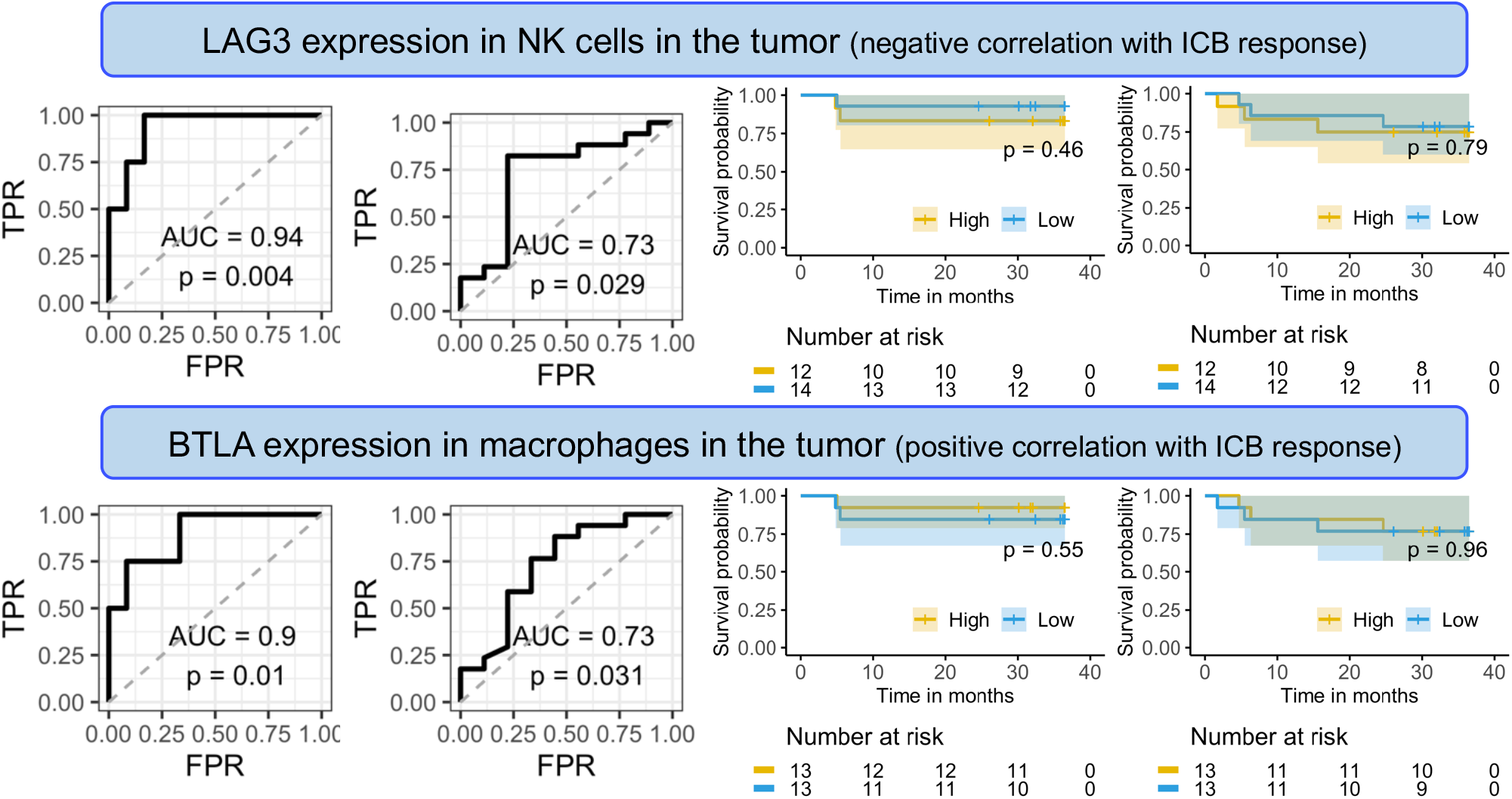
Two tumor ICG signatures are found to predict ICB response but patient survival after ICB treatment in HNSCC patients. The names of the two signatures are shown in the figure. The first two columns are ROC curves of ICB RECIST response and pathological response respectively. The last two columns are overall survival and progression-free survival after ICB treatment respectively. Note that the gene expression levels in the TME are predicted from the blood using the machine learning models. The data is from [31].

**Supplementary Figure 10.**
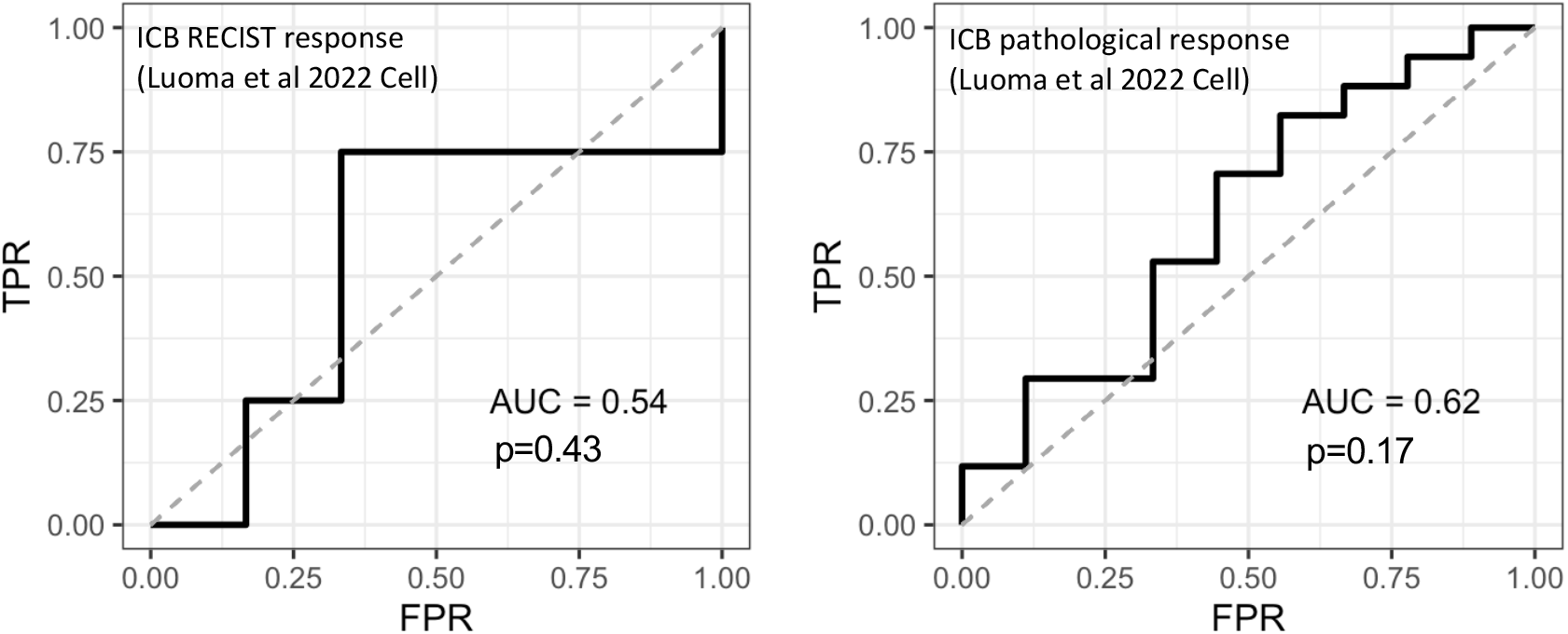
The ICFR signature of (B_memory_ -T_reg_) / (B_memory_ + T_reg_) is not found to predict ICB response in HNSCC patients when computed on the blood ICFs.

**Supplementary Figure 11.**
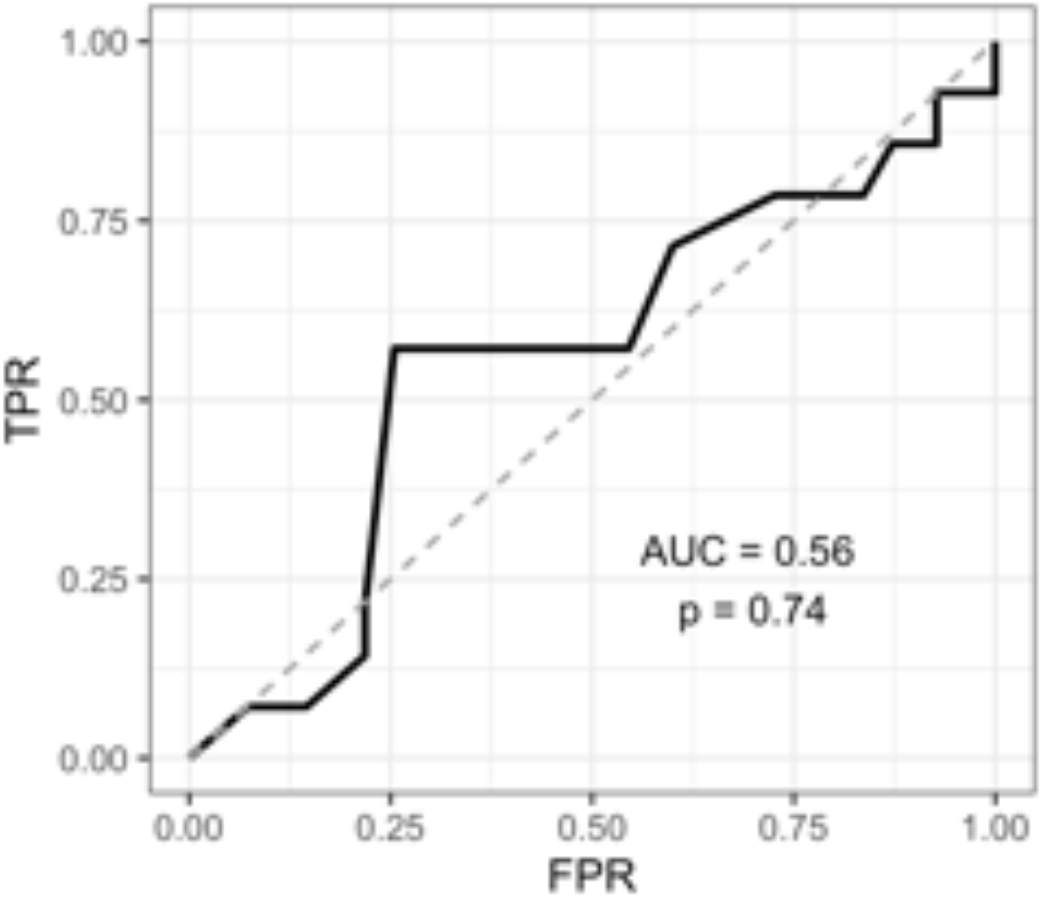
The predictive power of the TMB for ICB response in HNSCC patients in the MSK-IMPACT cohort. This cohort consists of in total 69 patients with 14 of them being responders as shown in data from [37].

**Supplementary Figure 12.**
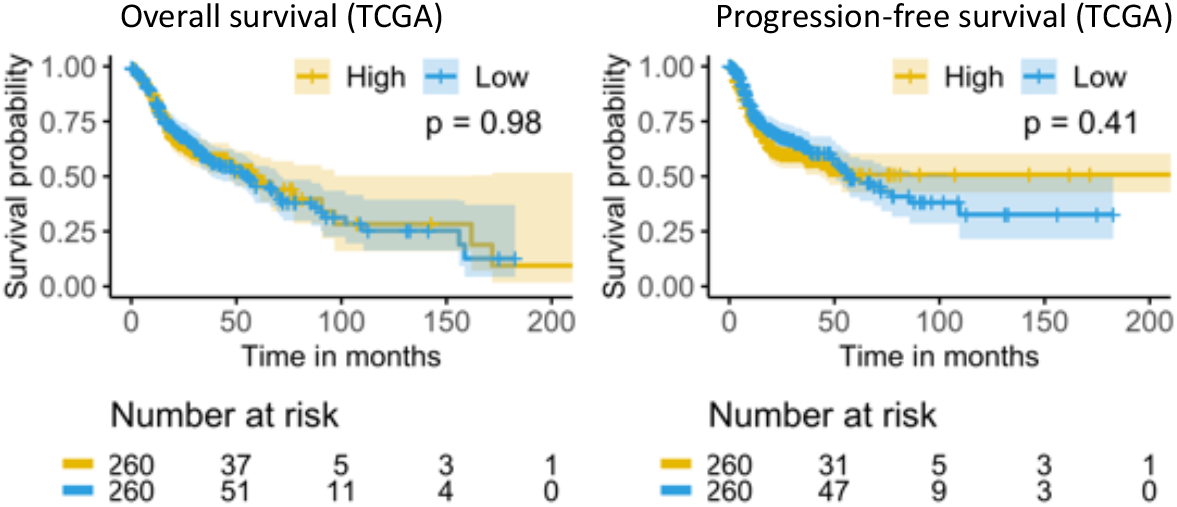
The tumor ICFR signature of (B_memory_ -T_reg_) / (B_memory_ + T_reg_) is not found to predict the survival of HNSCC patients in the TCGA dataset, where patients were not treated with ICB.

**Supplementary Figure 13.**
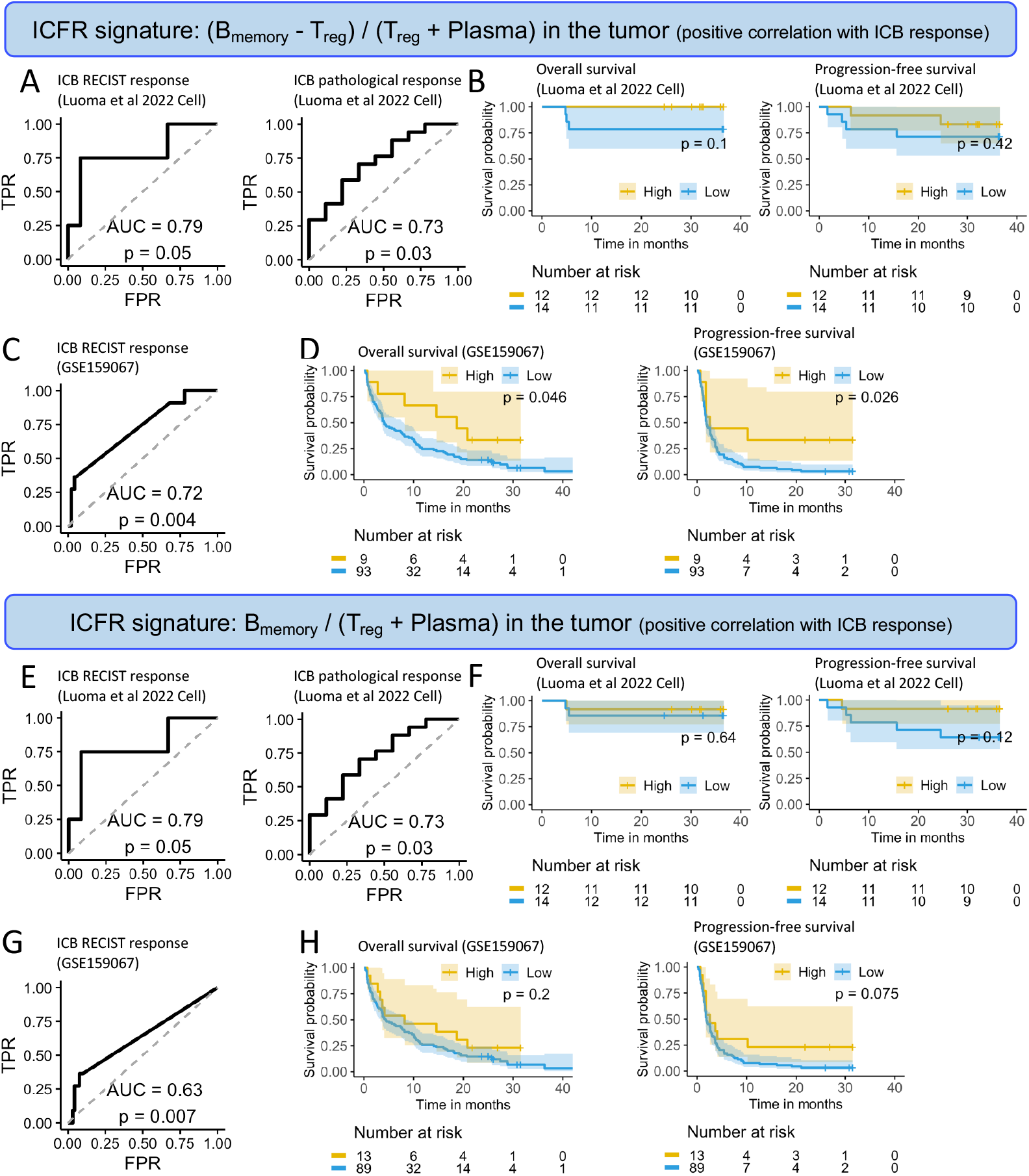
Two additional tumor ICFR signatures predict HNSCC patients’ ICB response but do not constantly predict patients’ survival after ICB treatment. The first ICFR signature is (B_memory_ - T_reg_) / (T_reg_ + Plasma) (**A-D)**, the second signature is B_memory_ / (T_reg_ + Plasma) (**E-H)**. ROC curves of ICB response prediction using the first ICFR signature on the single-cell dataset (**A**) and the bulk dataset (**C**). Survival analysis using the first ICFR signature on the single-cell dataset (**B**) and the bulk dataset (**D**). ROC curves of ICB response prediction using the second ICFR signature on the single-cell dataset (**E**) and the bulk dataset (**G**). Survival analysis using the second ICFR signature on the single-cell dataset (**F**) and the bulk dataset (**H**). It should be noted that tumor ICFRs on the single-cell dataset are predicted from the blood, while the tumor ICFRs on the bulk dataset are calculated from deconvolution results using CIBERSORT [36].

**Supplementary Material SM1:** Predictable genes in different immune cell types.

**Supplementary Material SM2:** Comprehensive lists of all signatures to predict patient ICB response.

